# Legal Harvest of Shorebirds and Resident Game Birds on a Caribbean island: A Martinique Case Study

**DOI:** 10.1101/2025.06.17.660260

**Authors:** Amelia R. Cox, Eduardo Gallo-Cajiao, Fred Tremblay, Fabian Rateau, Kévin Urvoy, Benoit Laliberté, Patrice Boniface, Dario Euphrosine, Tibo Lavanne, Christian Roy

## Abstract

Legal hunting practiced under regulatory frameworks is not always assessed for its sustainability. The lack of harvest data is of particular concern in the case of birds occurring on islands. Resident species on islands often exhibit limited geographic ranges and inherently small population sizes, rendering them particularly susceptible to environmental and anthropogenic threats. Islands also support seasonally fluctuating populations of migratory species, providing essential habitats for stopovers and non-breeding periods. Within this context, the sustainability of gamebird harvest practiced on islands within a regulatory framework needs to be evaluated to inform management and prevent negative impacts on populations. Here, we present a case study of an island-wide assessment of bird hunting estimates, based on data available from a legal hunting program, in Martinique, a tropical oceanic island part of the Caribbean biodiversity hotspot. Our objectives were to determine the magnitude of gamebird harvest, track changes in harvest levels over time, and assess harvest sustainability. Our assessment, which is based on hunting logbooks and hunter surveys between the 2012-2013 and 2021-2022 hunting seasons, provides harvest estimates for two groups of migratory species, shorebirds (*Charadriidae, Scolopacidae*) and teals (*Anatidae)*, and two groups of resident species (*Columbidae and Mimidae*). In Martinique, harvest management measures, combined with increased hunter’s engagement and enforcement of regulations, has led more stringent harvest regulations, improved compliance from hunters, and a reduction of Martinique’s national shorebird harvest. However, considering continued and rapid shorebirds population declines across the Western Atlantic flyway, shorebird harvest assessment at the flyway level remains a priority.

## 1. Introduction

Harvest assessments are an important tool to assess harvest sustainability and can be used to inform adaptive management and conservation strategies (Anderson et al., 2018; Holopainen et al., 2018). The importance of harvest assessments to inform harvest regulations aimed at sustainable resource use is well exemplified by some migratory gamebird hunting programs, such as those seen in waterfowl management in the U.S., Canada, and some European countries (Anderson et al., 2018; Hirschfeld et al., 2019; Holopainen et al., 2018). However, harvest data are not always available and when they are, data sometimes remain unassessed due to capacity issues (Hirschfeld et al., 2019). While managing populations without proper harvest assessment is possible, it limits the ability to evaluate management actions, achieve population goals, and prevent resource overexploitation (Blomberg et al., 2022; Eriksen et al., 2018; Weinbaum et al., 2013).

Overharvesting is a pervasive threat to animal and plant species (Díaz et al., 2019), yet quantitative harvest assessments at the species level are not always available, even when harvest is managed under a regulatory framework (Hirschfeld et al., 2019; Maxwell et al., 2016). The paucity of data on harvest is of particular concern in the case of birds who rely on island habitat as these species can be particularly vulnerable to harvest (Costello et al., 2013). Some island resident species have naturally restricted geographic ranges and consequently intrinsic small populations, which confer vulnerability (Matthews et al., 2022). Importantly, a high proportion (78%) of contemporary avian extinctions have occurred on oceanic islands with hunting having played a role in some cases (Szabo et al., 2012). Islands can also play an important role in the life cycle of migratory species as they can be used for stopovers during migration. While migratory species might only spend a short amount of time on islands, they can be exposed to harvest during their migration to threatening levels (McDuffie et al., 2022). Harvest management for migratory species can be particularly difficult since local harvest on a given island needs to be contextualized along the migration path and the cumulative harvest of the species across all jurisdictions can be difficult to estimate (Gallo-Cajiao et al., 2020; Watts and Turrin, 2016). For both resident species and migratory species, providing analytical tools to support harvest management is important to ensure hunting does not drive population declines or limit recovery efforts of threatened species (Eraud et al., 2021).

While introduced species and habitat loss have played an important role driving island population declines, hunting has also been documented as a contributing factor (Kirwan et al., 2019; Raffaele et al., 2020). Hunting of multiple bird species, both legally and illegally, has been practiced widely in the West Indies, and for various purposes such as subsistence, sport, and population control. Harvested species include both endemic species, such as parrots and pigeons, and migratory ones, such as shorebirds and ducks (Evans, 1991; Kirwan et al., 2019; Raffaele et al., 2020; Reed et al., 2018; Rivera-Milán et al., 2014; Wiley and Kirwan, 2013). Today, shorebird harvest remains a concern for population recovery in the Western Atlantic Flyway (AFSI Harvest Working Group, 2020; Watts et al., 2015).

Here, we present a case study of an island-wide assessment of game bird harvest in Martinique, a tropical island part of the Lesser Antilles archipelago in the West Indies, using data from a legal hunting monitoring program conducted by the Fédération des Chasseurs de Martinique (hereafter the Hunting Federation). The data collected has been used to inform adaptive harvest management actions over the years. Our aim is to help evaluate the sustainability of resident and migratory species harvest. Specifically, we ask the following questions: (i) what is the magnitude of harvest of bird species in Martinique as reported under a legal hunting framework; (ii) how has harvest changed over time, and (iii) how sustainable is the level of harvest. Based on hunting logbooks from the 2012-2013 to 2021-2022 hunting seasons, we provide annual harvest estimates for 6 resident species (4 Columbidae and 2 Mimidae), 9 migratory species (1 Charadriidae, 8 Scolopacidae) and one group (Anatidae), and evaluate the sustainability of migratory species harvest using a potential biological removal approach.

## 2. Methods

### 2.1 Study area

#### 2.1.1. Biophysical characteristics

Martinique is a tropical island in the eastern Caribbean Sea and the 3^rd^ largest island in the Lesser Antilles after Trinidad and Guadeloupe. With large differences in rainfall and topography across the island, habitats in Martinique are diverse, ranging from xerophytic shrublands and dry forest, to wet tropical forest, high altitude grasslands and savannas, and mangroves. A total of 183 bird species have been recorded in Martinique, of which about 74 breed on the island (BirdLife International, 2025).

Migratory bird species present in Martinique include several arctic and near-arctic breeding shorebirds which migrate through elliptical routes, meaning they migrate south through the Western Atlantic Flyway, but North through a more westerly route (Myers et al., 1990). Peak shorebird abundances in Martinique are typically recorded in September and October, overlapping with the hunting season. While some individuals will spend the rest of the non-breeding season in the region, most individuals will migrate further south. Resident species are mostly year-round residents in Martinique, although some species are presumably moving between Caribbean islands.

#### 2.1.2. Regulatory framework of bird hunting

In Martinique, authorized gamebird species are listed in the ministerial decree dated February 17, 1989, which includes 11 Anatidae, 10 Scolopacidae, 3 Charadriidae, 6 Colombidae and 2 Mimidae. Hunting regulations for each season are laid down in a yearly prefectural decree, which sets the opening and closing dates for each species, the days of the week when hunting is prohibited, daily or seasonal bag limits, etc. Decrees are established via a consultative process where the Direction de l’Environnement, de l’Aménagement et du Logement (DEAL) proposes a draft hunting decree which is then reviewed by the Commission Départementale de Gestion de la Chasse et de la Faune Sauvage (CDCFS). CDCFS is a hunting and wildlife management board, whose members are representatives of state agencies, such as DEAL and the Office Français de la Biodiviersité (OFB), the Hunting Federation and other hunting and conservation organizations. Most hunters in Martinique are members of one of the 34 local hunters’ associations which are coordinated by the departmental Hunting Federation. These associations lease land from the French State for hunting purposes, and only a handful of people hunt on non-public lands. After review by CDCFS, the prefect publishes the decree for a 20-day public consultation period. Once the consultation period is over, the DEAL reviews the comments received and publishes the yearly decree.

Game-bird regulations in Martinique have become considerably stricter over the last decade. Since 2013-2014, there have been multiple changes to harvest regulations for a range of species including changes to daily and seasonal bag limit and complete closure of harvest for some species.

### 2.2. Study design and data collection

#### 2.2.1. Annual harvest survey

Individuals who wish to renew their hunting license are required to fill out an annual harvest survey. In this survey, the Hunting Federation asks hunters to estimate their annual harvest from the previous hunting season, and the number of days spent hunting. Harvest estimates and efforts are reported either at the species level or the group level (e.g. harvest of all shorebird species is reported combined). This survey provides a baseline number of hunters who hunt or plan on hunting each year.

#### 2.2.2 Hunting Logbooks

When renewing their hunting license, hunters are also handed out a compulsory hunting logbook by the Hunting Federation. All hunters are required to carry an up-to-date logbook when in the field. In that logbook, they must register when and where they hunt and the number of birds from each species harvested that day (if any). Commonly harvested species are included in the logbook, including 10 shorebird species, 9 duck species and 8 resident species. However, the first version of the logbook did not include all harvested species and was updated in 2021-2022 to include additional species: black-bellied plover, ruddy turnstone, ring-billed duck, and lesser scaup.

Hunters typically diligently record dates of hunting activity especially when they are successful. These annual logbooks are collected by the Hunting Federation at the end of the season. Though hunters are obligated to maintain their logbooks up-to-date while hunting, many do not return the logbooks at the end of the hunting season; daily harvest for these hunters is therefore unknown. Hunter identities are not tracked in a way that allows for year-to-year comparison of logbooks, making it difficult to evaluate individual hunting success or link it to harvest outcomes.

### 2.3. Data analysis

#### 2.3.1. General model

Hunting activity often displays weekday/weekend cyclic patterns. To address this, we used the daily hunter activity and harvest data from the logbooks to calculate weekly activity (yes/no) and weekly harvest for each hunter throughout the hunting season. For the analysis, we used entries that were made during the legal hunting season only and we omitted all logbook entries for harvest before the season opened.

We modeled the harvest of 15 species and one group of species in Martinique. Although some harvest were reported for an additional 15 species, we excluded these species from our analysis due to insufficient harvest data (see Table S1 in Appendix A for a full list of species included in logbooks). For the species included, we modelled each species independently, except teals that we grouped together, using a hierarchical Bayesian model including three distinct components. We used generalized additive models for the first two components of the model: 1) weekly proportion of hunters actively harvesting the species and 2) weekly harvest per active hunter.

#### 2.3.2. Model for Migratory Species

We modeled the number of active hunters in week *w*, during hunting season *y*, that harvested the species in question using a binomial distribution such as:

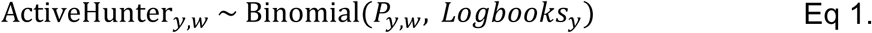

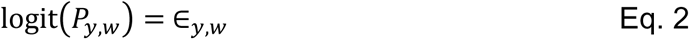

Where *P*_*y*,*w*_ is the proportion of hunters that were active in a given week out of the total number of returned logbooks. We used the “jagam” function of the *mgcv* R package (Wood, 2016) to set up basis functions to create a smooth for the week that varies by hunting season (k=9) which ∈_*y*,*w*_ is drawn from.

We modeled weekly harvest individually for each species using a truncated compound Gamma-Poisson distribution to account for over dispersion (i.e., variance larger than the mean; Greene, 2008) in weekly harvest as follows:

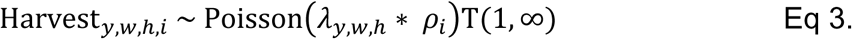

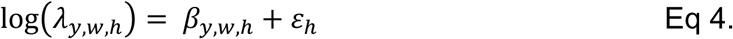

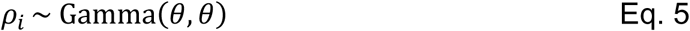

Where Harvest is the number of birds reported harvested at year *y*, during week *w*, by hunter *h*, during hunting day *i,* λ is the expected number of birds harvested, *ρ* is the overdispersion parameter. We truncated the distribution because the number of birds reported harvested is always either equal or greater than one. We modelled the expected numbers of birds on the log scale as a smooth term (β) that varies in function of the year, week and hunter. We created the basic function via the “jagam” function of the *mgcv* R package with k =9. We also included a random effect for the individual hunter h, as some hunters consistently harvest more than others. We drew the overdispersion terms from a Gamma distribution with the rate and shape set to a single parameter *θ*.

The final component of the model aimed at estimating the total harvest each year for a given species. The logbooks provide excellent information about harvest patterns for individual hunters within a season, but logbooks are not available for all hunters within a given year so summing up the harvest underestimates the total harvest. To alleviate this issue, in years where an annual survey was conducted, we used the total number of respondents to the survey to serve as a total hunter population. In years without an annual survey, we simulated the number of hunters based on the minimum and maximum number of hunters who filled the harvest survey over the length of our study (range from 574-881 hunters) and sampled the possible number of hunters from a uniform distribution. We used the number of annual survey respondents as a measure of the hunting population, rather than the number of hunting licenses issued, because many licensed individuals do not hunt and are solely licensed in order to legally own firearms. These individuals do not fill out the annual survey.

We then performed a data augmentation procedure, whereby we used the estimated proportion of hunters that did not return their logbook (e.g., Missing) for year *y*, that were active in week *w*.

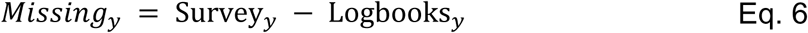

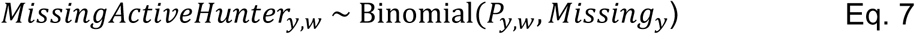

We then created a vector predicting missing harvest values for each missing hunter in each hunting season and week. We used a step function to prevent the model from creating more harvest values than necessary. We then estimated the total harvest by summing known logbook harvest with predicted harvest from the missing hunters, at both the seasonal and weekly level.

#### 2.3.3. Model for Resident Species

##### 2.3.3.1. Thrashers and Pigeons

We used a very similar approach to model harvest of thrashers and pigeons. However, since harvest was only allowed on weekends, starting October 1 for hunting seasons 2012-2013 through 2017-2018, we altered the model to take into account the changes in the regulations. We included a fixed effect to the active of the model to address the expected reduction in harvest as follow:

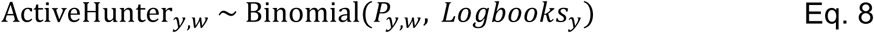

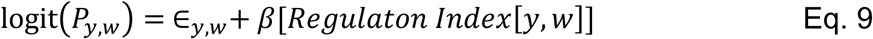

where *β*[*Regulation Index*[*y*, *w*]] is a term that accounts for changes in regulations. We also modified the total harvest model to account for the modifications in regulations:

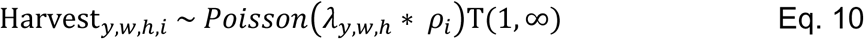

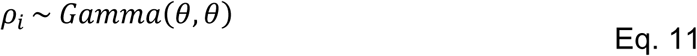

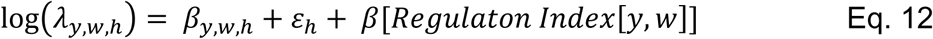

##### 2.3.3.2. Doves

For dove harvest, the short season (5 Sunday per year) required a simpler modeling approach. We therefore used a simpler model, with a fixed effect for week, and a random effect for harvest year in both the active hunter and weekly harvest components. Common ground dove harvest was low and could not support fixed effects for week so we modeled the annual mean.

#### 2.3.4. Model Estimation

All data processing was conducted in R version 4.3.1 (R Core Team, 2024). All analyses were conducted in JAGS version 4.3.0 (Plummer, 2003), called via the R package jagsUI (Kellner, 2021). We used non-informative priors for all parameters and ran six chains with randomized initial start values for 47,000 iterations, using the first 10,000 iterations as burn in and thinning the chains to save every 15th iteration, yielding 14,800 posterior samples for each parameter. We assessed model convergence visually and by making sure that the Gelman-Rubin statistic (*R*^^^) was less or equal than 1.1 (Gelman et al., 2013). We also used a posterior predictive check to evaluate the performance of each component of the model. All the JAGS models are included in Appendix B. We report medians with 90% highest density intervals calculated using the bayestestR package (Makowski et al., 2019)

#### 2.3.5. Derived Parameters

Potential Biological Removal (PBR) is an estimation of the maximum sustainable anthropogenic mortality and is based on a combination of population size and demographic rates of the species. Crucially, PBR incorporates anthropogenic mortality across the entire annual lifecycle and from all sources, not solely harvest. To assess sustainability of shorebird harvest in Martinique, we calculated what percentage of each species’ PBR the annual harvest estimates represents. In all cases we used the mean PBR reported in Watts et al. (2015).

## 3. Results

### 3.1. Annual harvest survey

Since 2015, a total of 3,607 hunters filled the annual harvest survey, with an average of 721 ± 114 hunters per season. According to the harvest survey results, the lowest number of active hunters was recorded in 2018-2019 with 574 hunters and the highest number of active hunters was in 2020-2021 with 881 hunters (Figure 2).

**Figure 1:**
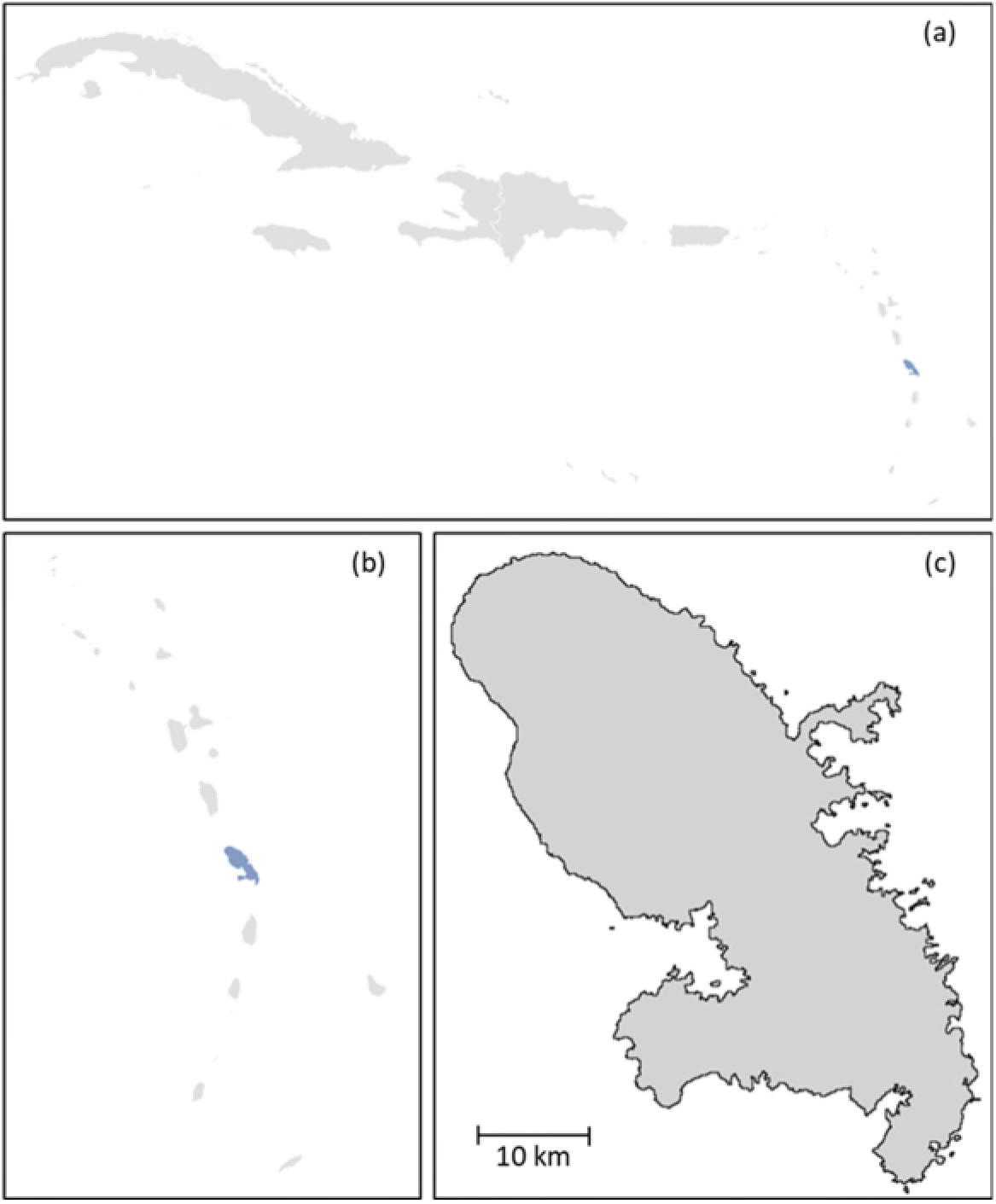
Map of Martinique at various scales (a: location of Martinique in the West Indies in light blue; b: location of Martinique in the Lesser Antilles in light blue; c: Martinique).

**Figure 2:**
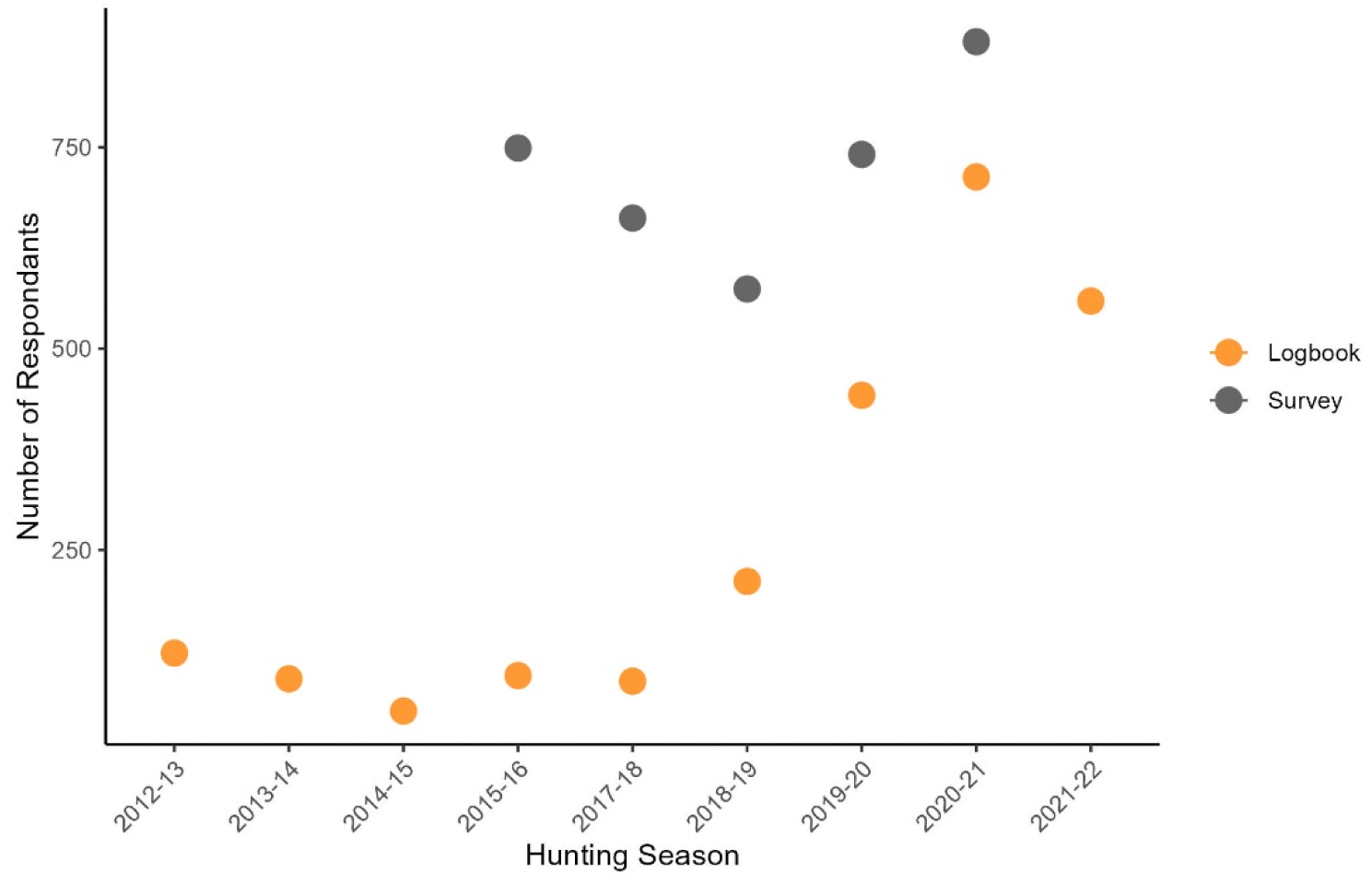
Number of hunters who filled the annual harvest survey (grey) and who submitted their hunting logbook (orange) at the end of each hunting season. The annual harvest survey began in 2015-2016.

Overall, 2,576 hunters submitted their logbooks over nine hunting seasons, out of which 301 logbooks did not contain any harvest data and were then excluded from analyses. The proportion of hunters submitting their logbooks increased significantly since 2015-2016, likely due to increased enforcement. Indeed, logbook return increased from 13% of registered hunters in 2015-2016 to 81% in 2020-2021.

### 3.2. Predicted Migratory Birds Harvest

#### 3.2.1 Shorebird Reported Harvest

Over nine hunting seasons, hunters collectively reported the harvest of 22 896 shorebirds and 4,470 teals in their logbooks., which yields an average of 2 544 [886 – 4,283] shorebirds and 497 [84 – 2,010] teals harvested per season.

#### 3.2.1. Shorebirds Modelled Harvest

Over the study period, our model estimated the average annual shorebird harvest at 6,984 individuals (Bayesian Credible intervals [BCI] 90% = 3,596 – 12,744). Harvest of combined shorebird species declined by 67% (BCI 90% = 61 - 73%) between 2012– 2013 and 2021–2022 hunting seasons. Mean annual harvest per hunter also declined during the survey period. There were, however, important yearly fluctuations, with the highest mean annual harvest occurring in 2015–2016 (18.8 shorebird/hunter; BCI 90% = 15.1 – 25.0) and the lowest in 2021–2022 (4.3 shorebird/hunter; BCI 90% = 4.0 – 4.7; Figure S1 – S26 Appendix C).

Mean harvest was highest for lesser yellowlegs (3,521; BCI 90% = 1,659 –9,249; Figure 3), followed by short-billed dowitcher (762; BCI 90% = 307 – 2,509), greater yellowlegs (715; BCI 90% =421 – 2,390) and American golden-plover (589; BCI 90% = 77 – 1,315; Figure 3; Table 1). When comparing the median annual harvest in Martinique to the mean PBR of the Western Atlantic Flyway (Watts et al. 2015), the local Martinique harvest represents 19.97% (BCI 90% = 4.85 – 48.47) of the Flyway’s PBR for whimbrel, 15.73% (BCI 90% = 6.34 – 51.77) for short-billed dowitcher, and 7.00% (BCI 90% = 4.12 – 23.41) for greater yellowlegs (Table 1).

**Figure 3.**
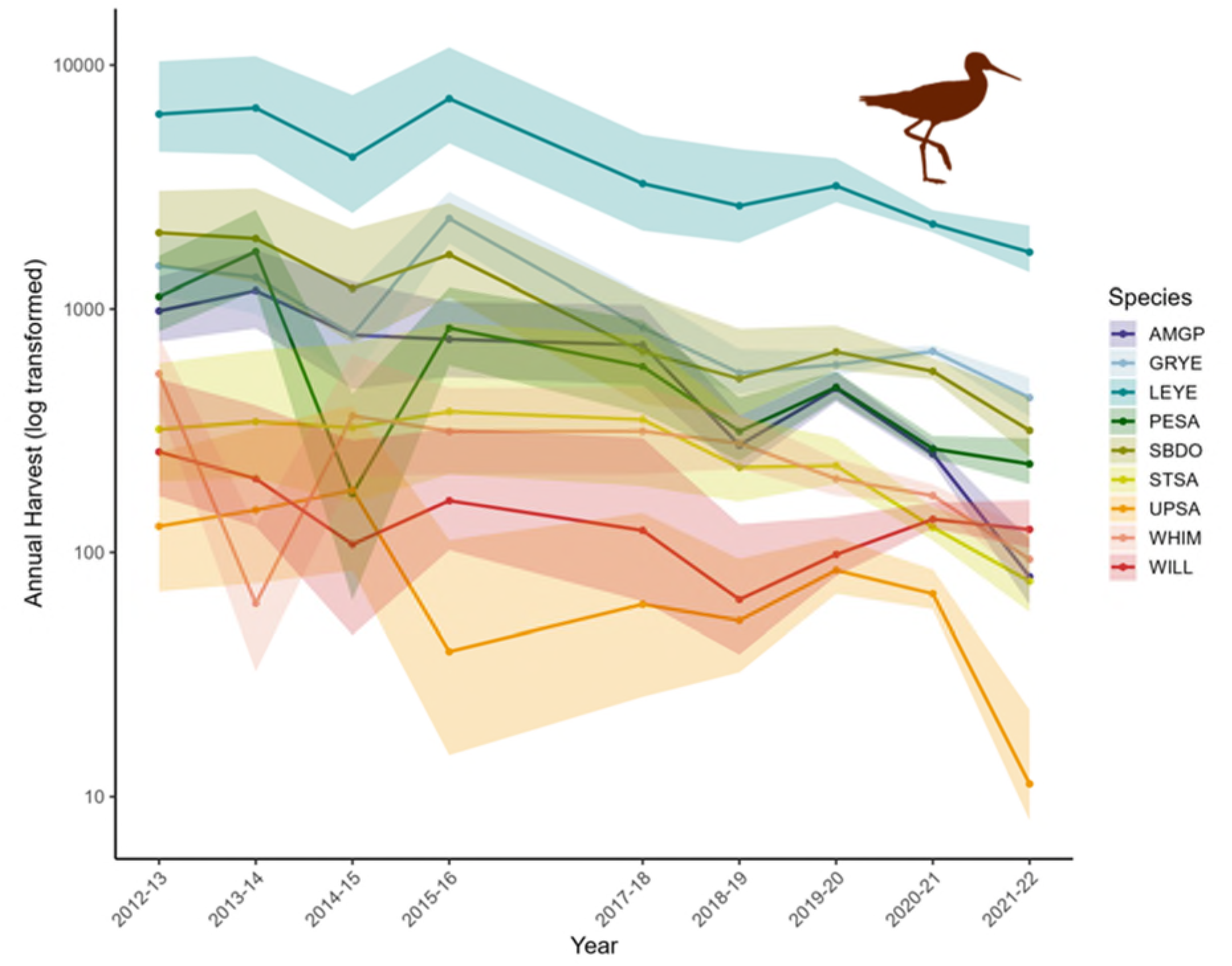
Annual harvest estimates for shorebird species harvested in Martinique during the legal hunting season between 2012-2013 and 2021-2022. Annual harvest estimates for shorebird species harvested in Martinique during the legal hunting season between 2012-2013 and 2021-2022 (AMGP: American golden plover; GRYE: greater yellowlegs; LEYE: lesser yellowlegs; PESA: pectoral sandpiper; SBDO: short-billed dowitcher; STSA: stilt sandpiper; UPSA: upland sandpiper; WHIM: whimbrel; WILL: willet). Solid lines represent median model predictions; the shaded areas represent the 90% Bayesian Credible Intervals.

**Table 1:**
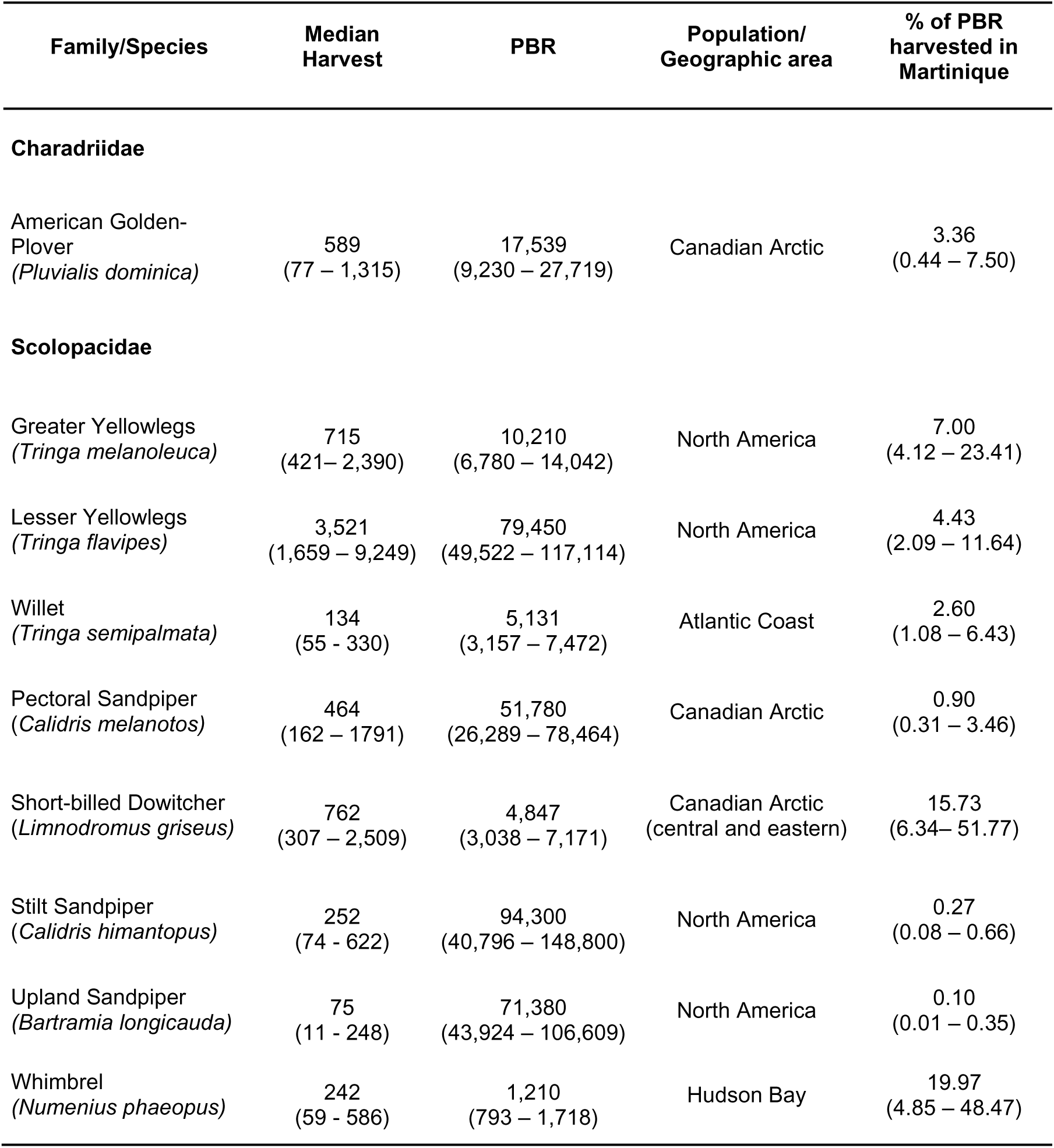
Medians and 90% Bayesian credible intervals for the annual harvest of shorebird species harvested in Martinique during the legal hunting season between 2012–2013 and 2021–2022 compared to their reported Potential Biological Removal (PBR; Watts et al. 2015).

| Family/Species | Median Harvest | PBR | Population/ Geographic area | % of PBR harvested in Martinique |
| --- | --- | --- | --- | --- |
| **Charadriidae** |  |  |  |  |
| American Golden-Plover<br>(*Pluvialis dominica*) | 589<br>(77 – 1,315) | 17,539<br>(9,230 – 27,719) | Canadian Arctic | 3.36<br>(0.44 – 7.50) |
| **Scolopacidae** |  |  |  |  |
| Greater Yellowlegs<br>(*Tringa melanoleuca*) | 715<br>(421– 2,390) | 10,210<br>(6,780 – 14,042) | North America | 7.00<br>(4.12 – 23.41) |
| Lesser Yellowlegs<br>(*Tringa flavipes*) | 3,521<br>(1,659 – 9,249) | 79,450<br>(49,522 – 117,114) | North America | 4.43<br>(2.09 – 11.64) |
| Willet<br>(*Tringa semipalmata*) | 134<br>(55 - 330) | 5,131<br>(3,157 – 7,472) | Atlantic Coast | 2.60<br>(1.08 – 6.43) |
| Pectoral Sandpiper<br>(*Calidris melanotos*) | 464<br>(162 – 1791) | 51,780<br>(26,289 – 78,464) | Canadian Arctic | 0.90<br>(0.31 – 3.46) |
| Short-billed Dowitcher<br>(*Limnodromus griseus*) | 762<br>(307 – 2,509) | 4,847<br>(3,038 – 7,171) | Canadian Arctic<br>(central and eastern) | 15.73<br>(6.34– 51.77) |
| Stilt Sandpiper<br>(*Calidris himantopus*) | 252<br>(74 - 622) | 94,300<br>(40,796 – 148,800) | North America | 0.27<br>(0.08 – 0.66) |
| Upland Sandpiper<br>(*Bartramia longicauda*) | 75<br>(11 - 248) | 71,380<br>(43,924 – 106,609) | North America | 0.10<br>(0.01 – 0.35) |
| Whimbrel<br>(*Numenius phaeopus*) | 242<br>(59 - 586) | 1,210<br>(793 – 1,718) | Hudson Bay | 19.97<br>(4.85 – 48.47) |

#### 3.2.2. Teals Modelled Harvest

On average, we estimate that a total of 1,075 (BCI 90% = 601 – 2,143) teals are harvested each season. According to the Hunting Federation, most of these are likely blue-winged teals as the occurrence of green-winged teal in Martinique during the hunting season is uncommon. Teal harvest varies considerably among hunting seasons. Hunting season 2018–2019 had the lowest harvest estimate of 682 (515 – 941) teals harvested and 2020–2021 the highest with 2,115 (2,077 – 2,171 Figure S16 – S17 Appendix C).

#### 3.2.3. Out-of-season harvest of migratory species

Of the 2576 logbooks we had access to, 21 reported shorebird harvest that was before the shorebird hunting season had started. Most of this harvest occurred in the 2012 – 2013 hunting season (32 of 37 dates), and mostly targeted lesser yellowlegs (203 of 309 shorebirds reported out of season).

### 3.3. Resident species harvest

Of the 2 576 logbooks collected, hunters reported 29,547 doves, 6,467 pigeons, and 1,333 thrashers harvested over nine hunting seasons, which yields an average of 867 [16 – 2,170] pigeons, 167 [11 – 628] thrashers, and 3 283 [704 – 11,993] doves harvested per season.

#### 3.3.1. Pigeon and Thrasher Modelled Harvest

On average, our model estimates that 2,174 (BCI 90% = 129 – 3,523) scaly-naped pigeons are harvested annually. However, pigeon harvest varied considerably over the years, with little harvest from 2013–2014 to 2015–2016, and up to 3,203 (BCI 90% = 2,109 – 5,674) individuals harvested in 2012-2013 and 2,740 (BCI 90% = 2,440 – 3,383) in 2019–2020.

Harvest of scaly-breasted and pearly-eyed thrashers is approximately 282 (59 – 958) annually (Figure 4). Harvest is split relatively evenly between the two species and their harvest levels have remained relatively unchanged over time (Figure S33 – S38 Appendix C).

**Figure 4:**
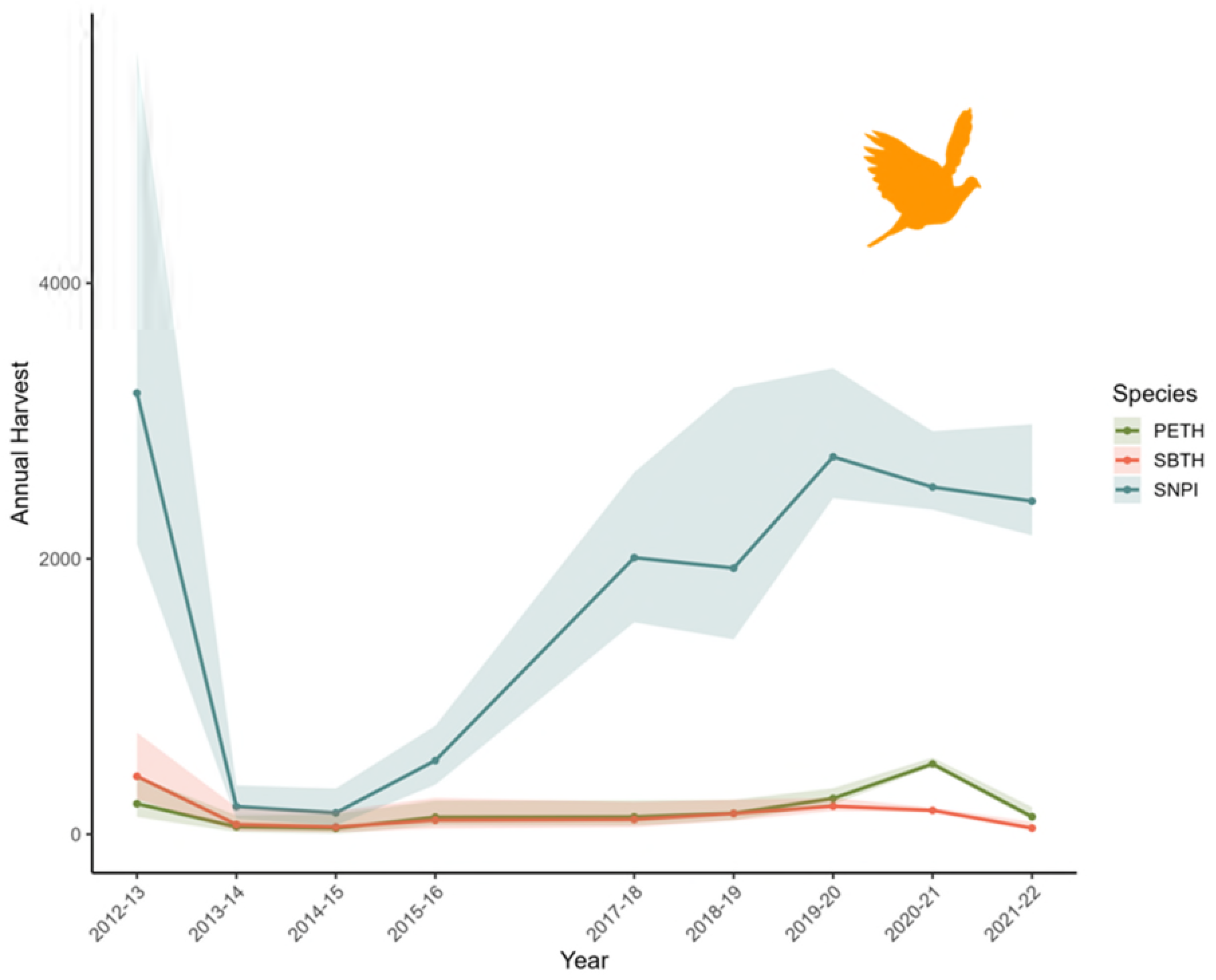
Annual harvest estimates for pigeon and thrashers species harvested in Martinique during the legal hunting season between 2012-2013 and 2021-2022 (PETH: pearly-eyed thrasher, SBTH: scaly-breasted thrasher, SNPI: scaly-naped pigeon). Solid lines represent median model predictions; the shaded areas represent the 90% Bayesian Credible Intervals

#### 3.3.2. Dove Modelled Harvest

Median annual harvest among resident doves was highest for zenaida dove (7,664BCI 90% = 2,453 – 12,135), followed by Eurasian collared dove (2,052; BCI 90% = 993 – 3,089; Table 2). Common ground dove harvest remained minimal throughout the years, with an average of 18 (BCI 90% = 0 – 208) individuals harvested each season where harvest was allowed. Hunting of common ground dove was closed for four hunting seasons between 2012–2013 and 2015–2016. While the annual harvest of the introduced Eurasian collared dove remained relatively stable, the reported harvest of zenaida dove varied considerably over the years (Figure 5; Figure S27 – S47 Appendix C).

**Figure 5:**
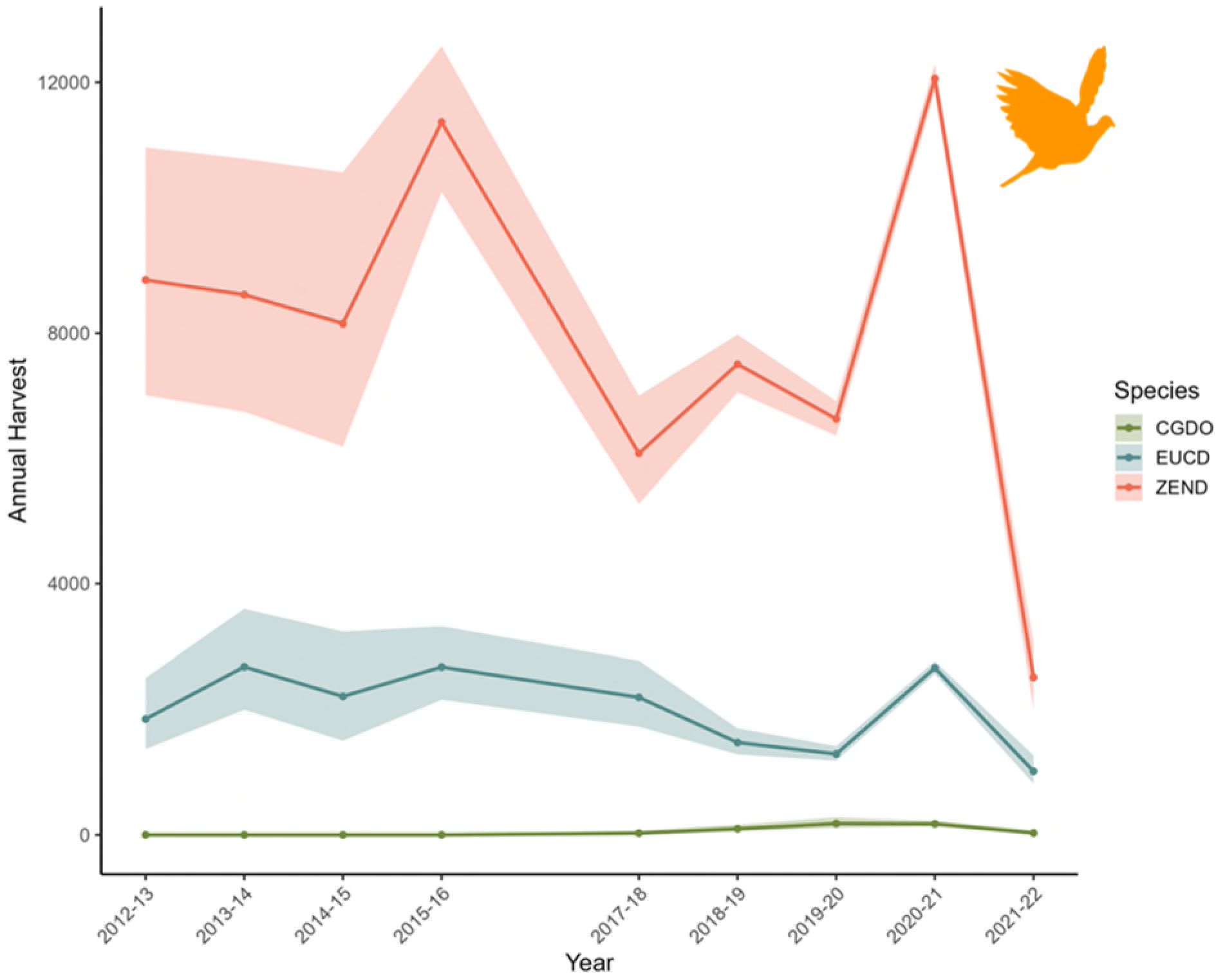
Annual harvest estimates for dove species harvested in Martinique during the legal hunting season between 2012-2013 and 2021-2022 (CGDO: common ground cove, EUCD: Eurasian collared dove, ZEND: zenaida dove). Solid lines represent median model predictions; the shaded areas represent the 90% Bayesian credible intervals.

**Table 2:**
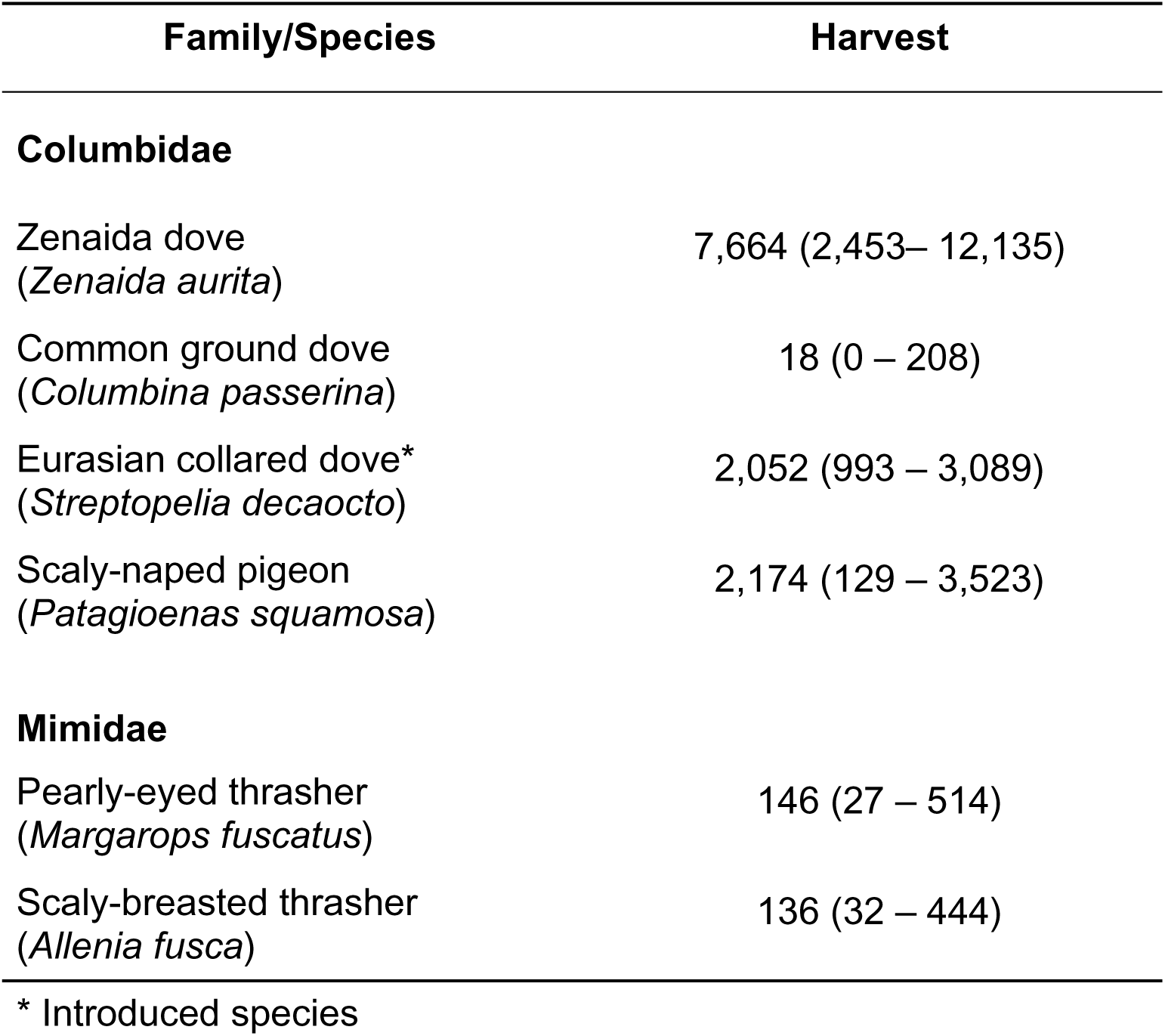
Medians and 90% Bayesian credible intervals for the mean harvest of doves, pigeons, and thrashers harvested in Martinique during the legal hunting season between 2012–2013 and 2021–2022. Median harvest for common ground dove reported only across years with a hunting season (2015–2016 to 2020–2021).

| Family/Species | Harvest |
| --- | --- |
| **Columbidae** |  |
| Zenaida dove<br>(*Zenaida aurita*) | 7,664 (2,453– 12,135) |
| Common ground dove<br>(*Columbina passerina*) | 18 (0 – 208) |
| Eurasian collared dove*<br>(*Streptopelia decaocto*) | 2,052 (993 – 3,089) |
| Scaly-naped pigeon<br>(*Patagioenas squamosa*) | 2,174 (129 – 3,523) |
| **Mimidae** |  |
| Pearly-eyed thrasher<br>(*Margarops fuscatus*) | 146 (27 – 514) |
| Scaly-breasted thrasher<br>(*Allenia fusca*) | 136 (32 – 444) |

\* Introduced species

#### 3.3.3 Out-of-season harvest of resident species

There were 36 instances of pigeon harvest outside the legal season, for a total of 72 scaly-naped pigeons and 3 white-crowned pigeons harvested. Of these 36 instances, 12 occurred before the opening of the hunting season and the remainder 24 occurred on weekdays during periods when hunting was only allowed on the weekend.

A total of 138 unique logbooks reported out-of-season harvest of Eurasian collared-dove, zenaida dove, or common ground dove. Of the 1674 doves that were reported to have been harvested out of season, 510 doves were harvested before the season started whereas the remaining 1,164 doves were harvested during the season but did not respect the “Sundays only” regulation.

## 4. Discussion

Our study advances game bird hunting assessment in Martinique by providing a modelling approach to obtain island-wide harvest estimates at the species level, and, in the case of shorebirds, comparing these to sustainable take levels at the flyway scale. Based on hunter logbooks, we successfully estimated the annual harvest for 15 species and one group of species legally hunted in Martinique over a nine-year period (2012– 2013 to 2021–2022). Between 2012–2013 and 2021–2022, annual shorebird harvest in Martinique declined by 67% (61%–73%). Harvest levels for resident species have been relatively stable, except zenaida and Eurasian-collared dove, for which harvest drastically declined in 2021–2022. Due to accelerating population declines observed in migratory shorebirds (Smith et al., 2023), conservation managers need to reconsider hunting as a contemporary threat, rather than a historical threat as has been commonly framed in conservation scholarship, policy, and management plans (Shrubb, 2013). Importantly, hunter engagement plays in developing and implementing management measures.

The decline in annual shorebird harvest observed after 2015–2016 season is likely driven by a combination of the implementation of more restrictive harvest regulations (i.e., daily and seasonal bag limits, season date restrictions and a moratorium on harvest for some species) and other external factors such as the COVID-19 pandemic during the 2021–2022 hunting season. Social pressure to reduce shorebird harvest levels increased after two radio-tracked Whimbrels were shot in 2011 on the neighboring island of Guadeloupe (All About Birds, 2011; Hopper, 2011). This pressure, combined with the subsequent creation of the Atlantic Shorebird Harvest Working Group under the Atlantic Flyway Shorebird Initiative (AFSI Harvest Working Group, 2017) played a role in the implementation of harvest restrictions across the Western Atlantic Flyway. As a result, the French State began adopting additional harvest restrictions on shorebird species for the 2013–2014 season and as of 2024–2025, management measures are now in place for 12 species of shorebirds, including five for which hunting is now closed (i.e. lesser yellowlegs, short-billed dowitcher, whimbrel, hudsonian godwit and ruddy turnstone).

Factors that led to the successful implementation of harvest management measures include the continuous engagement of hunters about shorebird population status and hunters’ participation on the harvest and wildlife management board. In parallel, the Hunting Federations played a key role by emphasizing the importance of returning diligently filled logbooks, which in turn provided insight on the current level of harvest. The information derived from this study contributed to the closure of harvest for species which PBR’s were proportionally the highest in Martinique (i.e. whimbrel and short-billed dowitcher) in 2022–2023, followed by closure of the most harvested species in term of absolute numbers (i.e. lesser yellowlegs) in 2024-2025.

Hunter engagement and education are important tools to ensure regulatory compliance (Blevins and Edwards, 2009), but the presence of law enforcement can also play a key role in ensuring adherence to the regulations (Jacoby, 2014; Von Essen et al., 2014). The importance of the perceived fairness of the regulatory-making process and the trust in the agencies in charge of the regulations can also play an important role in the acceptance of the regulations (Schroeder et al., 2017; Schroeder and Fulton, 2017; Von Essen et al., 2014). Adaptive management (AM) frameworks which engage hunters and stakeholders can be burdensome but are an effective way to secure stakeholder acceptance of harvest management decisions (Enck et al., 2006; Eriksson et al., 2024; Holmgaard et al., 2018; Williams and Madsen, 2013).

The changes in Martinique shorebird harvest regulations are promising. However, much work remains, especially when considering the cumulative impact of harvest along the entire flyway and the relatively unknown harvest levels in areas where harvest is suspected to be the highest (i.e., Suriname and Guyana; AFSI Harvest Working Group, 2020, 2017; Andres et al., 2022; Mizrahi et al., 2025). To fully understand the population-level impacts of shorebird harvest, obtaining harvest estimates throughout the Western Atlantic Flyway could be crucial to ensure proper harvest management and the perennity of the species throughout the flyway.

Harvest management of migratory shorebirds in Martinique presents intrinsic international dimensions. Managing hunting of migratory shorebirds is of utmost complexity as these species complete their life cycle across multiple political jurisdictions with a diversity of regulatory frameworks, capacity for enforcement, and value systems (Boardman 2006, Watts and Turrin 2016). In all, managing hunting of these species requires a flyway-wide approach exemplified by waterfowl hunting management in North America (Baldasarre and Bolen 1994). However, realizing such a goal across the 39 countries of the Western Hemisphere is very difficult with no mechanism currently in place for international coordination across the entire region. A first main hurdle to achieve sustainable hunting for migratory shorebirds is to have flyway-wide data on hunting magnitude, but detection of hunting and aggregation of data remain as two perennial issues. Even though these challenges largely remain to date, the 2016 Memorandum of Understanding signed by the US, Canada, and France, as it pertains to its overseas territories in the West Indies, intends to enable data sharing on shorebird ecology, conservation, and hunting management, providing some hope for achieving sustainable hunting.

Conversely, the conservation of resident gamebird species in Martinique relies largely on domestic policy. Of the six resident species hunted, one is introduced to the West Indies, two are distributed in the Neotropics beyond the West Indies, and three are endemic to the West Indies, of which one is endemic to the Lesser Antilles and none to Martinique. There are neither current estimates of what would constitute sustainable harvest, nor data on population estimates or trends for any these resident species. The Convention on Migratory Species does not apply to those species as they do not seem to cross international borders on a regular basis. The Protocol Concerning Specially Protected Areas and Wildlife to the Convention for the Protection and Development of the Marine Environment of the Caribbean Region, a regional international institutional arrangement with potential relevance, does not include any of such species under its annexes. Importantly, two species of pigeon (i.e., white-crowned pigeon, plain pigeon) occurring in the West Indies have been placed under this agreement’s appendices, conferring them protection. In addition to this particular agreement, the Convention on Biological Diversity, to which France is a party, has relevance as two of its core objectives are biodiversity conservation and sustainable use, but their actual operationalization in relation to hunting of those species remain uncertain. Additionally, local threats, such as the introduction of the small Indian mongoose (*Urva auropunctata*), has been identified as one of the main drivers of bird population declines, at least for some species (Louppe et al. 2021). This introduced mammal predates on the white-breasted thrasher (*Ramphocinclus brachyurus*; Gros-Desormeaux et al. 2015), a globally threatened species (EN, BirdLife International 2020), and so it has the potential to also affect the populations of the two species of thrashers subject to hunting. Until the conservation of these species is explicitly incorporated into international institutional arrangements, their conservation will rely solely, or at least primarily, on French domestic policy.

A current limitation of our harvest assessment is the lack of external data sources to complement our findings. While we successfully modeled harvests for commonly hunted species in Martinique, our model could not provide reliable estimates for species with sparse harvest data. For these less well-documented species, monitoring population changes through harvest data alone is not feasible. Additionally, the absence of robust population estimates complicates the assessment of long-term harvest sustainability for resident species or the effectiveness of season closures. For example, the scaly-naped pigeon (*Patagioenas squamosa*) population is considered to be declining across the West Indies (BirdLife International 2024) so it is difficult to assess the sustainability of the harvest in Martinique without a reliable population index for the population on the island.

When managing bird hunting in Martinique, consideration should be given not only to local extinction risk but also to ecosystem-based approaches as some species that are hunted, such as frugivores (e.g., scaly-naped pigeon, Perez-Rivera 1978), can play a key role in ecosystem function and structure. External data is also crucial for distinguishing the effects of regulations from other factors, such as stochastic events (Dalsgaard et al. 2007, Wiley and Wunderle 1993). For instance, extreme weather events like hurricanes can force shorebirds to land in large numbers in the Caribbean during their migration, potentially creating significant hunting opportunities and increasing harvest rates. Volcanic eruptions can have a detrimental effect, due to habitat loss and direct mortality, on the island’s bird resident populations (Joseph 2013). In all, the thresholds for sustainable hunting of birds in Martinique will depend on the demographic parameters of the species such as population size, adult survival and recruitment, all of which can be affected by anthropogenic threats and stochastic events. Enhancing our knowledge of these parameters could enhance the efficacy and success of the harvest management strategies.

## 5. Conclusions

The regulatory regime for hunting of migratory and resident bird species in Martinique has considerably improved since 2012–2013 in large part because of the development of a harvest regulation system that was implemented in conjunction with increased hunter’s engagement and regulation enforcement. Our study underscores the necessity of both domestic and international collaboration to address data gaps for informing legal hunting programs, which require not only data collection but also regular data analysis that feeds into the decision-making process in a timely fashion. A more systematic implementation of adaptive management at the local scale in Martinique, and throughout the Western Atlantic Flyway could bolster the conservation benefits of the current harvest management system (Eriksson et al., 2024; Holopainen et al., 2018). There will be some important challenges such a system would require the coordination of management and monitoring programs across multiples jurisdictions with diverse regulatory frameworks, capacity for enforcement, and value systems (Boardman, 2006; Watts and Turrin, 2016). Flyway-wide assessment of shorebird hunting and its sustainability has proven extremely difficult in the East Asian-Australasian Flyway (Gallo-Cajiao et al., 2020), and there is currently no mechanism or resources to coordinate harvest across the 39 countries of the Western Hemisphere despite repeated calls for such a system (Watts and Turrin, 2016). Nonetheless, a multi-lateral approach to management would provide the basis towards harvest sustainability.

## CRediT authorship contribution statement

**Amelia Cox:** Conceptualization, Methodology, Data curation, Formal analysis, Visualization; Writing - original draft, review & editing. **Eduardo Gallo-Cajia:** Writing - original draft, Writing - review and editing. **Fred Tremblay:** Data curation, Formal analysis, Visualization; Writing - review & editing. **Fabian Rateau:** Data curation, Writing - review & editing. **Kevin Urvoy:** Data curation. **Benoit Laliberté:** Writing - review & editing. **Partrice Boniface:** Investigation, Data curation. **Dario Euphorsine :** Investigation, Data curation. **Tibo Lavanne:** Investigation, Data curation. **Chistian Roy:** Conceptualization, Methodology, Formal analysis, Writing - review & editing.

## Funding

Funding was provided by Fédération Départementale des Chasseurs de la Martinique, Office Français de la Biodiversité, and Environment and Climate Change Canada.

## Declaration of competing interest

We declare that we do not have competing interests.

## Acknowledgements

Many thanks to the hunters from the Fédération Départementale des Chasseurs de la Martinique who have dutifully filled their hunting logbook and the annual harvest survey. Without them this research would not have been possible.

## Notes

### Competing Interest Statement

The authors have declared no competing interest.

## 6. Reference

AFSI Harvest Working Group, 2020. Actions for the Atlantic Flyway Shorebird Initiative’s Shorebird Harvest Working Group 2020–2025. U.S. Fish & Wildlife Service, Migratory Bird Program, Falls Church, VA, USA.

AFSI Harvest Working Group, 2017. Achieving a sustainable shorebird harvest in the Caribbean and northern South America, progress report, 2011–2017 (Unpublished report). U.S. Fish & Wildlife Service, Migratory Bird Program, Falls Church, VA, USA.

All About Birds, 2011. Second tracked Whimbrel dies in Guadeloupe [WWW Document]. URL https://www.allaboutbirds.org/news/second-tracked-whimbrel-dies-in-guadeloupe/ (accessed 2.18.25).

Anderson, M.G., Alisauskas, R.T., Batt, B.D., Blohm, R.J., Higgins, K.F., Perry, M.C., Ringelman, J.K., Sedinger, J.S., Serie, J.R., Sharp, D.E., others, 2018. The migratory bird treaty and a century of waterfowl conservation. The Journal of Wildlife Management 82, 247–259.

Andres, B.A., Moore, L., Cox, A.R., Frei, B., Roy, C., 2022. A preliminary assessment of shorebird harvest in coastal Guyana. Wader Study 129. 10.18194/ws.00263

Baldasarre, G.A., Bolen, E.G., 1994. Waterfowl ecology and management. New York: John Wiley and Sons, Inc.

BirdLife International, 2020. Species factsheet: Ramphocinclus brachyurus. Downloaded from http://datazone.birdlife.org/species/factsheet/white-breasted-thrasher-ramphocinclus-brachyurus

BirdLife International (2024). Species factsheet: Scaly-naped Pigeon Patagioenas squamosa. Downloaded from https://datazone.birdlife.org/species/factsheet/scaly-naped-pigeon-patagioenas-squamosa on 16/06/2025

BirdLife International, 2025. Country factsheet: Martinique (to France) [WWW Document]. URL https://datazone.birdlife.org/country/factsheet/martinique (accessed 2.18.25).

Blevins, K., Edwards, T., 2009. Wildlife crime, in: Miller, J.M. (Ed.), 21st Century Criminology: A Reference Handbook. Sage Publications, San Antonio, Texas, USA, pp. 557–564.

Blomberg, E.J., Ross, B.E., Cardinal, C.J., Ellis-Felege, S.N., Gibson, D., Monroe, A.P., Schwalenberg, P.K., 2022. Galliform exclusion from the Migratory Bird Treaty Act has produced an alternate conservation path, but no evidence for differences in population status. The Condor 124, duab051.

Boardman, R., 2006. The international politics of bird conservation: biodiversity, regionalism and global governance. Edward Elgar Publishing.

Costello, M.J., May, R.M., Stork, N.E., 2013. Can we name Earth’s species before they go extinct? science 339, 413–416.

Dalsgaard, B., Hilton, G.M., Gray, G.A.L., Aymer, L., Boatswain, J., Daley, J., et al., 2007. Impacts of a volcanic eruption on the forest bird community of Montserrat, Lesser Antilles. Ibis 149, 298–312. DOI: 10.1111/j.1474-919X.2006.00631.x.

Díaz, S., Settele, J., Brondízio, E., Ngo, H., Guèze, M., Agard, J., Arneth, A., Balvnera, P., Brauman, K., 2019. Summary for policymakers of the global assessment report on biodiversity and ecosystem services of the Intergovernmental Science-Policy Platform on Biodiversity and Ecosystem Services. IPBES Secretariat, Bonn, Germany.

Enck, J.W., Decker, D.J., Riley, S.J., Organ, J.F., Carpenter, L.H., Siemer, W.F., 2006. Integrating ecological and human dimensions in adaptive management of wildlife-related impacts. Wildlife Society Bulletin.

Eraud, C., Devaux, T., Villers, A., Johnson, F.A., Francesiaz, C., 2021. popharvest: An R package to assess the sustainability of harvesting regimes of bird populations. Ecology and Evolution 11, 16562–16571.

Eriksen, L.F., Moa, P.F., Nilsen, E.B., 2018. Quantifying risk of overharvest when implementation is uncertain. Journal of Applied Ecology 55, 482–493.

Eriksson, L., Johansson, M., Månsson, J., Sandström, C., Elmberg, J., 2024. Adaptive capacity in the multi-level management system of migratory waterbirds: A case study of participatory goose management in Sweden. Journal of Environmental Planning and Management 67, 522–541.

Evans, P.G., 1991. Status and conservation of Imperial and Red-necked Parrots Amazona imperalis and A. arausiaca on Dominica. Bird Conservation International 1, 11–32.

Gallo-Cajiao, E., Morrison, T.H., Woodworth, B.K., Lees, A.C., Naves, L.C., Yong, D.L., Choi, C.-Y., Mundkur, T., Bird, J., Jain, A., others, 2020. Extent and potential impact of hunting on migratory shorebirds in the Asia-Pacific. Biological Conservation 246, 108582.

Gelman, A., Carlin, J.B., Stern, H.S., Dunson, D.B., Vehtari, A., Rubin, D.B., 2013. Bayesian data analysis.

Greene, W., 2008. Functional forms for the negative binomial model for count data. Economics Letters 99, 585–590.

Gros-Desormeaux, J.R., Lesales, T., Tayalay, A.G. 2015. Behavioral observations on the White-breasted Thrasher (Ramphocinclus brachyurus brachyurus): conservation implications. Acta Ethologica 18, 197–208. DOI: 10.1007/s10211-014-0207-3.

Hirschfeld, A., Attard, G., Scott, L., 2019. Bird hunting in Europe. British Birds 112, 153– 166.

Holmgaard, S.B., Eythórsson, E., Tombre, I.M., 2018. Hunter opinions on the management of migratory geese: A case of stakeholder involvement in adaptive harvest management. Human Dimensions of Wildlife 23, 284–292.

Holopainen, S., Arzel, C., Elmberg, J., Fox, A.D., Guillemain, M., Gunnarsson, G., Nummi, P., Sjöberg, K., Väänänen, V.-M., Alhainen, M., Pöysä, H., 2018. Sustainable management of migratory European ducks: finding model species. Wildlife Biology. 10.2981/wlb.00336

Hopper, T., 2011. Machi the shorebird shot in Guadeloupe after surviving 6,000km hurricane odyssey [WWW Document]. National Post. URL https://nationalpost.com/news/canada/machi-the-shorebird-shot-in-guadeloupe-after-surviving-6000km-hurricane-odyssey (accessed 2.18.25).

Jacoby, K., 2014. Crimes against nature: Squatters, poachers, thieves, and the hidden history of American conservation. Univ of California Press.

Joseph, P., 2013. Mount Pele, an ecoclimate gradient generator. Landscape and the Environment 7, 27–41.

Kellner, K., 2021. jagsUI: A Wrapper Around ‘rjags’ to Streamline ‘JAGS’ Analyses. Kirwan, G.M., Levesque, A., Oberle, M., Sharpe, C.J., 2019. Bird of the West Indies, Lynx and BirdLife International Field Guide. Lynx Edicions, Barcelona.

Louppe, V., Herrel, A., Pisanu, B., Grouard, S., Veron, G., 2021. Assessing occupancy and activity of two invasive carnivores in two Caribbean islands: implications for insular ecosystems. Journal of Zoology 313, 182–194. DOI: 10.1111/jzo.12845.

Makowski, D., Ben-Shachar, M.S., Lüdecke, D., 2019. bayestestR: Describing effects and their uncertainty, existence and significance within the Bayesian framework. Journal of Open Source Software 4, 1541.

Matthews, T.J., Wayman, J.P., Cardoso, P., Sayol, F., Hume, J.P., Ulrich, W., Tobias, J.A., Soares, F.C., Thébaud, C., Martin, T.E., others, 2022. Threatened and extinct island endemic birds of the world: Distribution, threats and functional diversity. Journal of Biogeography 49, 1920–1940.

Maxwell, S.L., Fuller, R.A., Brooks, T.M., Watson, J.E., 2016. Biodiversity: The ravages of guns, nets and bulldozers. Nature 536, 143–145.

McDuffie, L.A., Christie, K.S., Harrison, A.-L., Taylor, A.R., Andres, B.A., Laliberté, B., Johnson, J.A., 2022. Eastern-breeding Lesser Yellowlegs are more likely than western-breeding birds to visit areas with high shorebird hunting during southward migration. Ornithological Applications 124, duab061. 10.1093/ornithapp/duab061

Mizrahi, D., Spaans, A.L., Cox, A.R., Laliberté, B., Gallo-Cajiao, E., Roy, C., 2025. A national assessment of waterbird hunting in coastal wetlands of Suriname.

Myers, J., Sallaberry A, M., Ortiz, E., Castro, G., Gordon, L., Maron, J.L., Schick, C., Tabilo, E., Antas, P., Below, T., 1990. Migration routes of new world sanderlings (Calidris alba). The Auk 107, 172–180.

Perez-Rivera, R. A. 1978. Preliminary work of the feeding habits, nesting habitat and reproductive activities of the plain pigeon (Columba inornata wetmorei) and the red-necked pigeon (Columba squamosa), sympatric species: an analysis of their interaction. Science-Ciencia 5:89–98.

Plummer, M., 2003. JAGS: A program for analysis of Bayesian graphical models using Gibbs sampling. Proceedings of the 3rd International Workshop on Distributed Statistical Computing (DSC 2003).

R Core Team, 2024. R: A Language and Environment for Statistical Computing. R Foundation for Statistical Computing, Vienna, Austria.

Raffaele, H.A., Wiley, J., Garrido, O.H., Keith, A., Raffaele, J.I., 2020. Birds of the West Indies Second Edition. Princeton University Press.

Reed, E.T., Kardynal, K.J., Horrocks, J.A., Hobson, K.A., 2018. Shorebird hunting in Barbados: Using stable isotopes to link the harvest at a migratory stopover site with sources of production. The Condor 120, 357–370. 10.1650/CONDOR-17-127.1

Rivera-Milán, F.F., Boomer, G.S., Martínez, A.J., 2014. Monitoring and modeling of population dynamics for the harvest management of scaly-naped pigeons in Puerto Rico. The Journal of wildlife management 78, 513–521.

Schroeder, S.A., Fulton, D.C., 2017. Voice, perceived fairness, agency trust, and acceptance of management decisions among Minnesota anglers. Society & Natural Resources 30, 569–584.

Schroeder, S.A., Fulton, D.C., Lawrence, J.S., Cordts, S.D., 2017. How hunter perceptions of wildlife regulations, agency trust, and satisfaction affect attitudes about duck bag limits. Human Dimensions of Wildlife 22, 454–475.

Shrubb, M., 2013. Feasting, fowling and feathers: A history of the exploitation of wild birds. A&C Black.

Smith, P.A., Smith, A.C., Andres, B., Francis, C.M., Harrington, B., Friis, C., Morrison, R.G., Paquet, J., Winn, B., Brown, S., 2023. Accelerating declines of North America’s shorebirds signal the need for urgent conservation action. Ornithological Applications 125, duad003.

Szabo, J.K., Khwaja, N., Garnett, S.T., Butchart, S.H., 2012. Global patterns and drivers of avian extinctions at the species and subspecies level.

Von Essen, E., Hansen, H.P., Nordström Källström, H., Peterson, M.N., Peterson, T.R., 2014. Deconstructing the poaching phenomenon: A review of typologies for understanding illegal hunting. British Journal of Criminology 54, 632–651.

Watts, B., Reed, E., Turrin, C., 2015. Estimating sustainable mortality limits for shorebirds using the Western Atlantic Flyway. Wader Study. 10.18194/ws.00005

Watts, B.D., Turrin, C., 2016. Assessing hunting policies for migratory shorebirds throughout the Western Hemisphere. Wader Study 123, 6–15.

Weinbaum, K.Z., Brashares, J.S., Golden, C.D., Getz, W.M., 2013. Searching for sustainability: are assessments of wildlife harvests behind the times? Ecology letters 16, 99–111.

Wiley, J.W., Wunderle, J.M., 1993. The effects of hurricanes on birds, with special reference to Caribbean islands. Bird Conservation International 3, 319–349. DOI: 10.1017/S0959270900002598.

Wiley, J.W., Kirwan, G.M., 2013. The extinct macaws of the West Indies, with special reference to Cuban Macaw Ara tricolor. Bulletin of the British Ornithologists’ Club 133, 125–156.

Williams, J.H., Madsen, J., 2013. Stakeholder perspectives and values when setting waterbird population targets: Implications for flyway management planning in a European context. PloS one 8, e81836.

Wood, S.N., 2016. Just another Gibbs additive modeler: interfacing JAGS and mgcv. Journal of Statistical Software 75, 1–15.

