## Appendix A for "Legal Harvest of Shorebirds and Resident Game Birds on a Caribbean island: A Martinique Case Study"

**Table S1:** List of species included in harvest logbooks. Asterix denotes species that were included in the analyses.

| **Common Name** | **Local Name** | **Latin Name** | **Status** |
|---|---|---|---|
| Greater yellowlegs* | Clin | *Tringa melanoleuca* | Migrant |
| Willet* | Ailes blanches | *Tringa semipalmata* | Migrant |
| Lesser yellowlegs* | Patte jaune | *Tringa flavipes* | Migrant |
| Hudsonian godwit | Barge à queue noire | *Limosa haemastica* | Migrant |
| Short-billed dowitcher* | Grise à long bec | *Limnodromus griseus* | Migrant |
| Whimbrel* | Bec crochu | *Numenius phaeopus* | Migrant |
| Pectoral Sandpiper* | Dos rouge | *Calidris melanotos* | Migrant |
| Wilson’s snipe | Bécassine | *Gallinago delicata* | Migrant |
| American golden plover* | Pluvier doré | *Pluvialis dominica* | Migrant |
| Ruddy turnstone | Pluvier des salines | *Arenaria interpres* | Migrant |
| Upland sanpiper* | Poule vergene | *Bartramia longicauda* | Migrant |
| Stilt Sandpiper* | Chevalier à pieds verts | *Calidris himantopus* | Migrant |
| Black-bellied plover | Pluvier argenté | *Pluvialis squatarola* | Migrant |
| Blue-winged teal* | Sarcelle à ailes bleue | *Spatula discors* | Migrant |
| Green-winged teal* | Sarcelle à ailes vertes | *Anas crecca* | Migrant |
| American wigeon | Canard siffleur | *Mareca americana* | Migrant |
| American black duck | Canard noir | *Anas rubripes* | Migrant |
| Ring-billed duck | Fulligule à collier | *Aythya collaris* | Migrant |
| Lesser scaup | Petit fulligule | *Aythya affinis* | Migrant |
| Northern shoveler | Canard souchet | *Spatula clypeata* | Migrant |
| Mallard | Colvert | *Anas platyrhynchos* | Migrant |
| Black-bellied whistling duck | Dendrocygne à ventre noire | *Dendrocygna autumnalis* | Migrant |
| Fulvous whislting duck | Dendrocygne fauve | *Dendrocygna bicolor* | Migrant |
| Northern pintail | Canard pilet | *Anas acuta* | Migrant |
| Zenaida dove* | Tourterelle à queue carré | *Zenaida aurita* | Resident |
| Eurasian collared dove* | Tourterelle turque | *Streptopelia decaocto* | Resident |
| Scaly-naped pigeon* | Ramier cou rouge | *Patagioenas squamosa* | Resident |
| White-crowned pigeon | Ramier à tête blanche | *Patagioenas leucocephala* | Resident |
| Scaly-breasted thrasher* | Grive fine | *Allenia fusca* | Resident |
| Pearly-eyed thrasher* | Grosse grive | *Margarops fuscatus* | Resident |
| Common-ground dove* | Ortolan | *Columbina passerina* | Resident |
| Eared dove | Tourterelle oreillarde | *Zenaida auriculata* | Resident |

**Table S2:** Harvest regulation since 2012 for species included our analyses.

| | 2012-2013 | 2013-2014 | 2014-2015 | 2015-2016 to 2019-2020 | | 2020-2021 | | 2021-2022 | | 2022-2023 | 2023-2024 |
|---|---|---|---|---|---|---|---|---|---|---|---|
| **Species** | Daily | Daily | Daily | Daily | Seasonal | Daily | Seasonal | Daily | Seasonal | Daily | Daily |
| Whimbrel | | | 5 | 3 | 15 | 2 | 10 | 2 | 10 | | |
| American golden plover | | | | | | | | 12 | 50 | 5 combined with BBPL | 4 |
| Short-billed dowitcher | | | | | | | | 15 | | 5* | |
| Lesser yellowleg | | | | | | | | 20 | | 5 | 5 |
| Greater yellowleg | | | | | | | | | | | 10 |
| Willet | | | | | | | | | | | 10 |
| Stilt sandpiper | | | | | | | | | | | 10 |
| Pectoral sandpiper | | | | | | | | | | | 10 |
| Upland sandpiper | | | | | | | | | | | 10 |
| Shorebirds sp. | | | | | | | | | | 20 | 20 |
| Blue-winged teal | | | | | | | | | | | |
| Green-wing teal | | | | | | | | | | | |
| Scaly-naped pigeon | | | | | | | | | | 10 | 10 |
| Pearly-eyed thrasher | | | | | | | | 15 | | 3 | 5 |
| Scaly-breasted thrasher | | | | | | | | | | | 10 |
| Common ground dove | | | | 3 | 15 | 3 | 15 | | | | |
| Zenaida dove | | | | | | | | | | | |
| Eurasian collared dove | | | | | | | | | | | |

Moratorium on hunting    No bag limit
