## Appendix B for "Legal Harvest of Shorebirds and Resident Game Birds on a Caribbean island: A Martinique Case Study"

**Appendix B:** JAGS harvest models

### 2.1 GAM for shorebird harvest

```
model {
#For shorebirds and teal
########################################
#Active Hunters in a given week (out of total hunters)
#"Active" is defined as successfully harvesting the species
  eta <- X %*% b ## linear predictor

for (i in 1:Nobs) {
	logit(mu[i]) <-  eta[i]
	ActiveHunter[i] ~ dbin(mu[i],TotalHunter_logbook[YrIndex[i]])
	rate_hunt[YrIndex[i], WkIndex[i]] <- mu[i]
 } ## response

 ## Parametric effect priors CHECK tau=1/11^2 is appropriate!
  for (i in 1:1) { b[i] ~ dnorm(0,0.0087) }
  ## prior for s(Week)...
  K1 <- S1[1:9,1:9] * lambda[1]  + S1[1:9,10:18] * lambda[2]
  b[2:10] ~ dmnorm(zero[2:10],K1)
  ## prior for s(Week,Hunting.Season)...
  for (i in c(11:18,21:28,31:38,41:48,51:58,61:68,71:78,81:88, 91:98)) { b[i] ~ dnorm(0, lambda[3]) }
  for (i in c(19,29,39,49,59,69,79,89, 99)) { b[i] ~ dnorm(0, lambda[4]) }
  for (i in c(20,30,40,50,60,70,80,90, 100)) { b[i] ~ dnorm(0, lambda[5]) }
  ## smoothing parameter priors CHECK...
  for (i in 1:5) {
    lambda[i] ~ dgamma(.05,.005)
    rho[i] <- log(lambda[i])
  }


#########################################
#Total harvest of the given species per week for each individual.
#Total harvest-1 is modelled b/c 0's were removed previously (i.e. those individuals didn't hunt)
 theta_total ~ dgamma(1,0.1)
 eta_total <- X_total %*% b_total ## linear predictor
  for (i in 1:Nobs_t) {
	log(l_total[i]) <-  eta_total[i]
	rho_nbtotal[i] ~ dgamma( theta_total, theta_total)
	Total[i] ~ dpois(l_total[i]* rho_nbtotal[i]) T(0,)
	  } ## response
  ## Parametric effect priors CHECK tau=1/15^2 is appropriate!
  for (i in 1:1) { b_total[i] ~ dnorm(0,0.0046) }
  ## prior for s(Week)...
  K1_total <- S1_total[1:9,1:9] * lambda_total[1]  + S1_total[1:9,10:18] * lambda_total[2]
  b_total[2:10] ~ dmnorm(zero_total[2:10],K1_total)
  ## prior for s(Week,Hunting.Season)...
  for (i in c(11:18,21:28,31:38,41:48,51:58,61:68,71:78,81:88, 91:98)) { b_total[i] ~ dnorm(0,
lambda_total[3]) }
  for (i in c(19,29,39,49,59,69,79,89, 99)) { b_total[i] ~ dnorm(0, lambda_total[4]) }
  for (i in c(20,30,40,50,60,70,80,90, 100)) { b_total[i] ~ dnorm(0, lambda_total[5]) }
  ## prior for s(Logbook.ID)...
  for (i in c(101:dim_log)) { b_total[i] ~ dnorm(0, lambda_total[6]) }
  ## smoothing parameter priors CHECK...
```

```
  for (i in 1:6) {
    lambda_total[i] ~ dgamma(.05,.005)
    rho_total[i] <- log(lambda_total[i])
  }
#Mean harvest (i.e. remove the specific

mu_eta_total <- X %*% b_total[1:100]

for (i in 1:Nobs) {
            log(mu_l_total[i]) <-  mu_eta_total[i]
            mu_lambda_total[YrIndex[i], WkIndex[i]] <- mu_l_total[i]
 }
#######################################################

#Derived values
#For each year that that we have survey data in
#addition to the logbooks, we should know exactly
#how many hunters there are that DIDN't fill in their logbooks
for (t in 4:8){ #Only have survey data for subset of the years.
for(w in 1:Nweek){
            ActiveHunter_survey[t, w] ~ dbin(rate_hunt[t, w] , (TotalHunter_survey[t-3]-
TotalHunter_logbook[t]))
            TotalActiveHunter[t,w] <- ActiveHunter_survey[t,w] + ActiveHunter_logbook[t,w]
}
TotalPerHunter[t] <- TotalAnnualHarvest[t]/TotalHunter_survey[t-3]
}


for (t in 1:3){
T_unknown[t] ~ dunif(Min, Max)
for(w in 1:Nweek){
            ActiveHunter_survey[t, w] ~ dbin(rate_hunt[t, w] , (round(T_unknown[t])-TotalHunter_logbook[t]))
            TotalActiveHunter[t,w] <- ActiveHunter_survey[t,w] + ActiveHunter_logbook[t,w]
}
TotalPerHunter[t] <- TotalAnnualHarvest[t]/T_unknown[t]
}

#In 2022, survey wasn't administered to everyone
T_unknown[9] ~ dunif(559,Max)
for(w in 1:Nweek){
            ActiveHunter_survey[9, w] ~ dbin(rate_hunt[9, w] , (round(T_unknown[9])-
TotalHunter_logbook[9]))
            TotalActiveHunter[9,w] <- ActiveHunter_survey[9,w] + ActiveHunter_logbook[9,w]
}
TotalPerHunter[9] <- TotalAnnualHarvest[9]/T_unknown[9]

for (t in 1:Nyear){
for(w in 1:Nweek){
Total_survey[t,w] <- mu_lambda_total[t, w] * ActiveHunter_survey[t,w]
TotalHarvest[t,w] <- TotalHarvest_logbook[t,w] + Total_survey[t,w]
}

TotalAnnualHarvest[t] <- sum(TotalHarvest[t,])
TotalAnnualDaysHunting[t] <- sum(TotalActiveHunter[t,])

}
```

```
##################################################
#Fit statistics
for (i in 1:Nobs) {
		ld_hunt[i] <- logdensity.bin(ActiveHunter[i] , mu[i],TotalHunter_logbook[YrIndex[i]])
		ActiveHunter_new[i] ~ dbin(mu[i],TotalHunter_logbook[YrIndex[i]])
		ld_hunt_new[i] <- logdensity.bin(ActiveHunter_new[i] , mu[i],TotalHunter_logbook[YrIndex[i]])
 }
  for (i in 1:Nobs_t) {
		ld_total[i] <- logdensity.pois (Total[i] , l_total[i]*rho_nbtotal[i])
		Total_new[i] ~ dpois(l_total[i] * rho_nbtotal[i]) T(0,)
		ld_total_new[i] <- logdensity.pois (Total_new[i] , l_total[i]*rho_nbtotal[i])
  }

fit_hunt <- sum(-2* ld_hunt)
fit_hunt_new <- sum(-2*ld_hunt_new)

fit_total <- sum(-2*ld_total)
fit_total_new <- sum(-2*ld_total_new)
}
```

### 2.2 GAM for pigeon and thrashers harvest

```
model {
#For shorebirds and teal
#######################################
#Active Hunters in a given week (out of total hunters)
#"Active" is defined as successfully harvesting the species
  eta <- X %*% b ## linear predictor

for(r in 1:2){ #To account for regulation changes from weekdays to weekends only
beta[r] ~ dnorm(0, pow(5, -2))
}


for (i in 1:Nobs) {
            logit(mu[i]) <-  eta[i] + beta[RegIndex[i]]
            ActiveHunter[i] ~ dbin(mu[i],TotalHunter_logbook[YrIndex[i]])
            rate_hunt[YrIndex[i], WkIndex[i]] <- mu[i]
 } ## response

 ## Parametric effect priors CHECK tau=1/11^2 is appropriate!
  for (i in 1:1) { b[i] ~ dnorm(0,0.0087) }
  ## prior for s(Week)...
  K1 <- S1[1:9,1:9] * lambda[1]  + S1[1:9,10:18] * lambda[2]
  b[2:10] ~ dmnorm(zero[2:10],K1)
  ## prior for s(Week,Hunting.Season)...
  for (i in c(11:18,21:28,31:38,41:48,51:58,61:68,71:78,81:88, 91:98)) { b[i] ~ dnorm(0, lambda[3]) }
  for (i in c(19,29,39,49,59,69,79,89, 99)) { b[i] ~ dnorm(0, lambda[4]) }
  for (i in c(20,30,40,50,60,70,80,90, 100)) { b[i] ~ dnorm(0, lambda[5]) }
  ## smoothing parameter priors CHECK...
  for (i in 1:5) {
    lambda[i] ~ dgamma(.05,.005)
    rho[i] <- log(lambda[i])
  }


########################################
#Total harvest of the given species per week for each individual.
#Total harvest-1 is modelled b/c 0's were removed previously (i.e. those individuals didn't hunt)
 theta_total ~ dgamma(1,0.1)
 eta_total <- X_total %*% b_total ## linear predictor
  for (i in 1:Nobs_t) {
            log(l_total[i]) <-  eta_total[i] + beta_total[RegIndexTot[i]]
            rho_nbtotal[i] ~ dgamma( theta_total, theta_total)
            Total[i] ~ dpois(l_total[i]* rho_nbtotal[i]) T(0,)

  } ## response

for(r in 1:2){beta_total[r] ~ dnorm(0,pow(5,-2))}

  ## Parametric effect priors CHECK tau=1/15^2 is appropriate!
  for (i in 1:1) { b_total[i] ~ dnorm(0,0.0046) }
  ## prior for s(Week)...
  K1_total <- S1_total[1:9,1:9] * lambda_total[1]  + S1_total[1:9,10:18] * lambda_total[2]
  b_total[2:10] ~ dmnorm(zero_total[2:10],K1_total)
  ## prior for s(Week,Hunting.Season)...
```

```
  for (i in c(11:18,21:28,31:38,41:48,51:58,61:68,71:78,81:88, 91:98)) { b_total[i] ~ dnorm(0,
lambda_total[3]) }
  for (i in c(19,29,39,49,59,69,79,89, 99)) { b_total[i] ~ dnorm(0, lambda_total[4]) }
  for (i in c(20,30,40,50,60,70,80,90, 100)) { b_total[i] ~ dnorm(0, lambda_total[5]) }
  ## prior for s(Logbook.ID)...
  for (i in c(101:dim_log)) { b_total[i] ~ dnorm(0, lambda_total[6]) }
  ## smoothing parameter priors CHECK...
  for (i in 1:6) {
    lambda_total[i] ~ dgamma(.05,.005)
    rho_total[i] <- log(lambda_total[i])
  }
#Mean harvest (i.e. remove the specific

mu_eta_total <- X %*% b_total[1:100]

for (i in 1:Nobs) {
        log(mu_l_total[i]) <-  mu_eta_total[i] + beta_total[RegIndex[i]]
        mu_lambda_total[YrIndex[i], WkIndex[i]] <- mu_l_total[i]
 }
#########################################################

#Derived values
#For each year that that we have survey data in
#addition to the logbooks, we should know exactly
#how many hunters there are that DIDN't fill in their logbooks
for (t in 4:8){ #Only have survey data for subset of the years.
for(w in 1:Nweek){
        ActiveHunter_survey[t, w] ~ dbin(rate_hunt[t, w] , (TotalHunter_survey[t-3]-
TotalHunter_logbook[t]))
        TotalActiveHunter[t,w] <- ActiveHunter_survey[t,w] + ActiveHunter_logbook[t,w]
}
TotalPerHunter[t] <- TotalAnnualHarvest[t]/TotalHunter_survey[t-3]
}


for (t in 1:3){
T_unknown[t] ~ dunif(Min, Max)
for(w in 1:Nweek){
        ActiveHunter_survey[t, w] ~ dbin(rate_hunt[t, w] , (round(T_unknown[t])-TotalHunter_logbook[t]))
        TotalActiveHunter[t,w] <- ActiveHunter_survey[t,w] + ActiveHunter_logbook[t,w]
}
TotalPerHunter[t] <- TotalAnnualHarvest[t]/T_unknown[t]
}

#In 2022, survey wasn't administered to everyone
T_unknown[9] ~ dunif(559,Max)
for(w in 1:Nweek){
        ActiveHunter_survey[9, w] ~ dbin(rate_hunt[9, w] , (round(T_unknown[9])-
TotalHunter_logbook[9]))
        TotalActiveHunter[9,w] <- ActiveHunter_survey[9,w] + ActiveHunter_logbook[9,w]
}
TotalPerHunter[9] <- TotalAnnualHarvest[9]/T_unknown[9]


for (t in 1:Nyear){
for(w in 1:Nweek){
```

```
		Total_survey[t,w] <- mu_lambda_total[t, w] * ActiveHunter_survey[t,w]
		TotalHarvest[t,w] <- TotalHarvest_logbook[t,w] + Total_survey[t,w]
}

TotalAnnualHarvest[t] <- sum(TotalHarvest[t,])
TotalAnnualDaysHunting[t] <- sum(TotalActiveHunter[t,])

}

####################################################
#Fit statistics
for (i in 1:Nobs) {
		ld_hunt[i] <- logdensity.bin(ActiveHunter[i] , mu[i],TotalHunter_logbook[YrIndex[i]])
		ActiveHunter_new[i] ~ dbin(mu[i],TotalHunter_logbook[YrIndex[i]])
		ld_hunt_new[i] <- logdensity.bin(ActiveHunter_new[i] , mu[i],TotalHunter_logbook[YrIndex[i]])
 }
  for (i in 1:Nobs_t) {
		ld_total[i] <- logdensity.pois (Total[i] , l_total[i]*rho_nbtotal[i])
		Total_new[i] ~ dpois(l_total[i] * rho_nbtotal[i]) T(0,)
		ld_total_new[i] <- logdensity.pois (Total_new[i] , l_total[i]*rho_nbtotal[i])
  }

fit_hunt <- sum(-2* ld_hunt)
fit_hunt_new <- sum(-2*ld_hunt_new)

fit_total <- sum(-2*ld_total)
fit_total_new <- sum(-2*ld_total_new)
}
```

### 2.3 Model for dove harvest

```
model {
#For generic doves (which is actually just EADO and EUCD)
########################################
#Active Hunters in a given week (out of total hunters)
#"Active" is defined as successfully harvesting the species
for (i in 1:Nobs) {
		ActiveHunter[i] ~ dbin(rate_hunt[YrIndex[i], WkIndex[i]],TotalHunter_logbook[YrIndex[i]])
 } ## response

#Priors on hunting activity
mu_hunt  ~ dnorm(0,pow(5,-2))
sigma.year ~ dt(0,pow(1,-2),1)T(0,)		#Cauchy scale set to 1
sigma.week ~ dt(0,pow(1,-2),1)T(0,)
for (w in 1:Nweek){
		epsilon_week[w] ~ dnorm(0,pow(sigma.week,-2))
}

for (t in 1:Nyear){
				epsilon_year[t] ~ dnorm(0, pow(sigma.year, -2))
}

for(w in 1:Nweek){
for(t in 1:Nyear){
				eta[w,t] <- mu_hunt + epsilon_year[t] + epsilon_week[w]
		logit(rate_hunt[t,w]) <- eta[w,t]
```

```
}}


##########################################
#Total harvest of the given species per week for each individual.
#Total harvest-1 is modelled b/c 0's were removed previously (i.e. those individuals didn't hunt)
 theta_total ~ dgamma(1,0.1)
   for (i in 1:Nobs_t) {
        log(l_total[i]) <- log_mu_lambda_total + epsilon_harv_year[YrtotIndex[i]] #+
epsilon_harv_week[WktotIndex[i]] # + epsilon.hunter[HunterIndex[i]]
        rho_nbtotal[i] ~ dgamma( theta_total, theta_total)
        Total[i] ~ dpois(l_total[i]* rho_nbtotal[i]) T(0,)

  } ## response


#Priors on harvest of active hunters.
#Priors on hunting activity
log_mu_lambda_total ~ dnorm(0,pow(5,-2))


sigmaharv.year ~ dt(0,pow(1,-2),1)T(0,)      #Cauchy scale set to 1

for (t in 1:Nyear){
        epsilon_harv_year[t] ~ dnorm(0, pow(sigmaharv.year, -2))
        log(mu_lambda_total[t])<- log_mu_lambda_total + epsilon_harv_year[t]
}




###########################################################

#Derived values
#For each year that that we have survey data in
#addition to the logbooks, we should know exactly
#how many hunters there are that DIDN't fill in their logbooks
for (t in 4:8){ #Only have survey data for subset of the years.
for(w in 1:Nweek){
        ActiveHunter_survey[t, w] ~ dbin(rate_hunt[t, w] , (TotalHunter_survey[t-3]-
TotalHunter_logbook[t]))
        TotalActiveHunter[t,w] <- ActiveHunter_survey[t,w] + ActiveHunter_logbook[t,w]
}
TotalPerHunter[t] <- TotalAnnualHarvest[t]/TotalHunter_survey[t-3]
}


for (t in 1:3){
T_unknown[t] ~ dunif(Min, Max)
for(w in 1:Nweek){
        ActiveHunter_survey[t, w] ~ dbin(rate_hunt[t, w] , (round(T_unknown[t])-TotalHunter_logbook[t]))
        TotalActiveHunter[t,w] <- ActiveHunter_survey[t,w] + ActiveHunter_logbook[t,w]
}
TotalPerHunter[t] <- TotalAnnualHarvest[t]/T_unknown[t]
}
T_unknown[9] ~ dunif(559,Max)
```

```
for(w in 1:Nweek){
	ActiveHunter_survey[9, w] ~ dbin(rate_hunt[9, w] , (round(T_unknown[9])-
TotalHunter_logbook[9]))
	TotalActiveHunter[9,w] <- ActiveHunter_survey[9,w] + ActiveHunter_logbook[9,w]
}
TotalPerHunter[9] <- TotalAnnualHarvest[9]/T_unknown[9]

for (t in 1:Nyear){
for(w in 1:Nweek){
Total_survey[t,w] <- mu_lambda_total[t] * ActiveHunter_survey[t,w]
TotalHarvest[t,w] <- TotalHarvest_logbook[t,w] + Total_survey[t,w]
}

TotalAnnualHarvest[t] <- sum(TotalHarvest[t,])
TotalAnnualDaysHunting[t] <- sum(TotalActiveHunter[t,])

}


###################################################
#Fit statistics
for (i in 1:Nobs) {
	ld_hunt[i] <- logdensity.bin(ActiveHunter[i] , rate_hunt[YrIndex[i],
WkIndex[i]],TotalHunter_logbook[YrIndex[i]])
	ActiveHunter_new[i] ~ dbin(rate_hunt[YrIndex[i], WkIndex[i]],TotalHunter_logbook[YrIndex[i]])
	ld_hunt_new[i] <- logdensity.bin(ActiveHunter_new[i] , rate_hunt[YrIndex[i],
WkIndex[i]],TotalHunter_logbook[YrIndex[i]])
 }
  for (i in 1:Nobs_t) {
	ld_total[i] <- logdensity.pois (Total[i] , l_total[i]*rho_nbtotal[i])
	Total_new[i] ~ dpois(l_total[i] * rho_nbtotal[i]) T(0,)
	ld_total_new[i] <- logdensity.pois (Total_new[i] , l_total[i]*rho_nbtotal[i])
  }

fit_hunt <- sum(-2* ld_hunt)
fit_hunt_new <- sum(-2*ld_hunt_new)

fit_total <- sum(-2*ld_total)
fit_total_new <- sum(-2*ld_total_new)
}
```
