## Appendix C for "Legal Harvest of Shorebirds and Resident Game Birds on a Caribbean island: A Martinique Case Study"

**Appendix C:** Additional Figures

**Migratory Species**

*American Golden Plover*

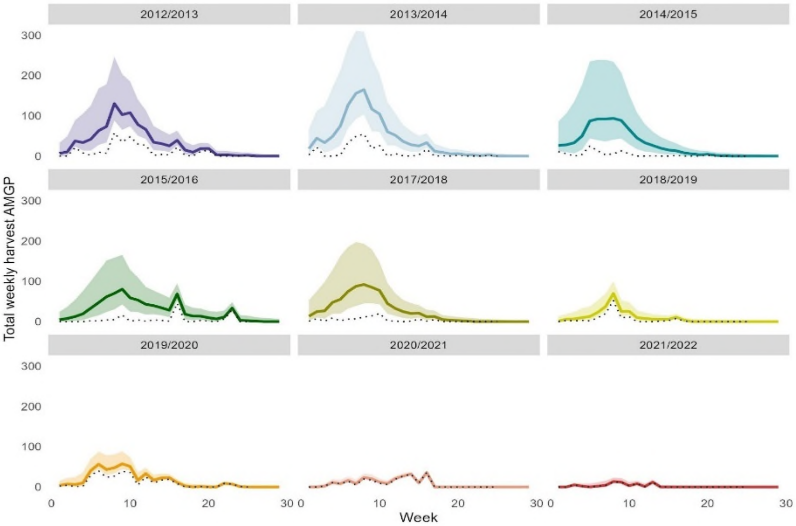


**Figure S1:** Estimated weekly harvest of American golden plover. The solid lines represent weekly national harvest using median model predicted weekly harvest, with a shaded area to represent the 90% BCIs on weekly harvest. The dotted line represents the weekly harvest reported in the logbooks.

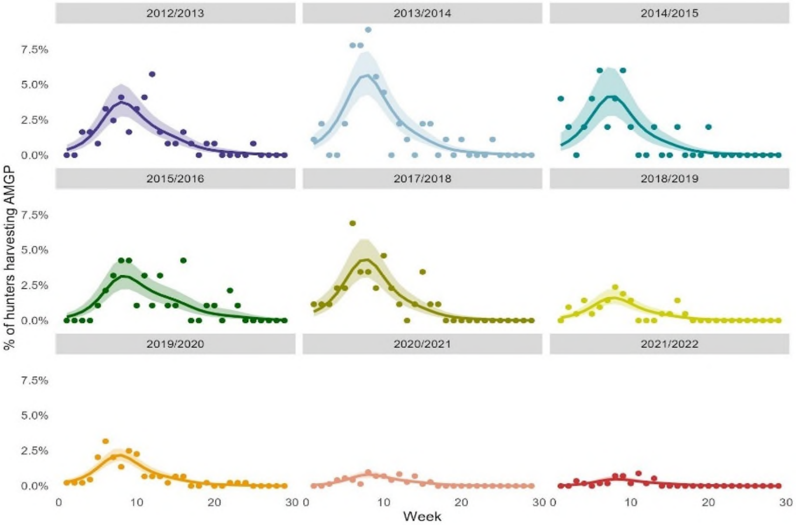


**Figure S2:** Estimated proportion of hunters that successfully harvested American golden plover per week. Solid lines represented the median model predicted proportion of hunters harvesting American golden plovers. The shaded areas represent the 90% BCIs. Points represent the percentage of logbooks in a given week that report harvest of American golden plovers.

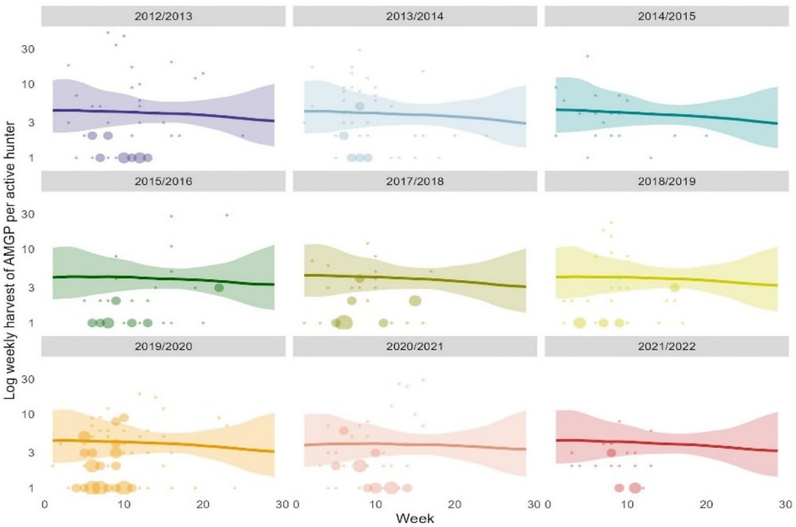


**Figure S3:** Reported weekly harvest of American golden plover per successful hunter. Solid lines represented median model predicted harvest per hunter. The shaded areas represent the 90% BCIs. Points represent logbook reported harvest, with point size representing the number of hunters reporting the given weekly harvest.

*Greater yellowlegs*

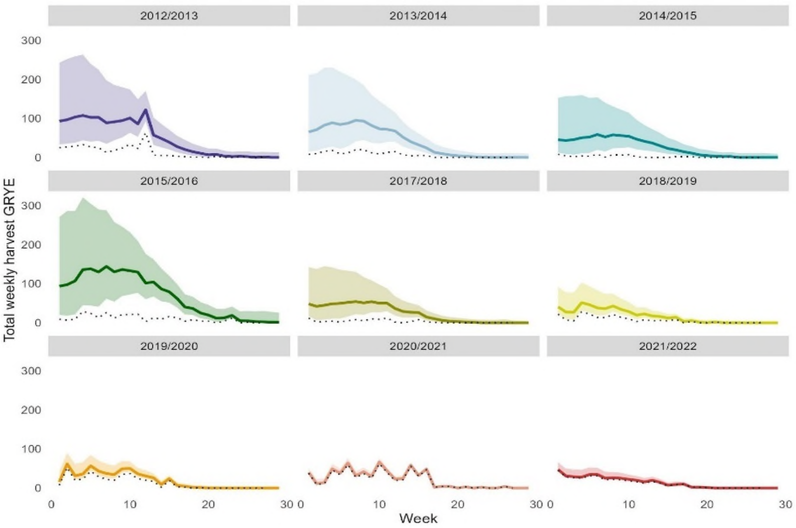


**Figure S4:** Estimated weekly harvest of Greater yellowlegs. The solid lines represent weekly national harvest using median model predicted weekly harvest, with a shaded area to represent the 90% BCIs on weekly harvest.The dotted line represents the weekly harvest reported in the logbooks.

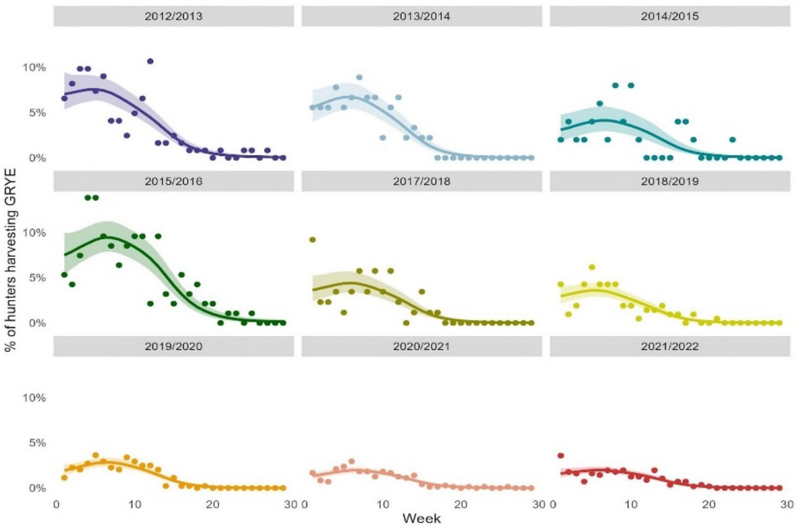


**Figure S5:** Estimated proportion of hunters that successfully harvested Greater yellowlegs per week. Solid lines represented the median model predicted proportion of hunters harvesting Greater yellowlegs. The shaded areas represent the 90% BCIs. Points represent the percentage of logbooks in a given week that report harvest of Greater yellowlegs.

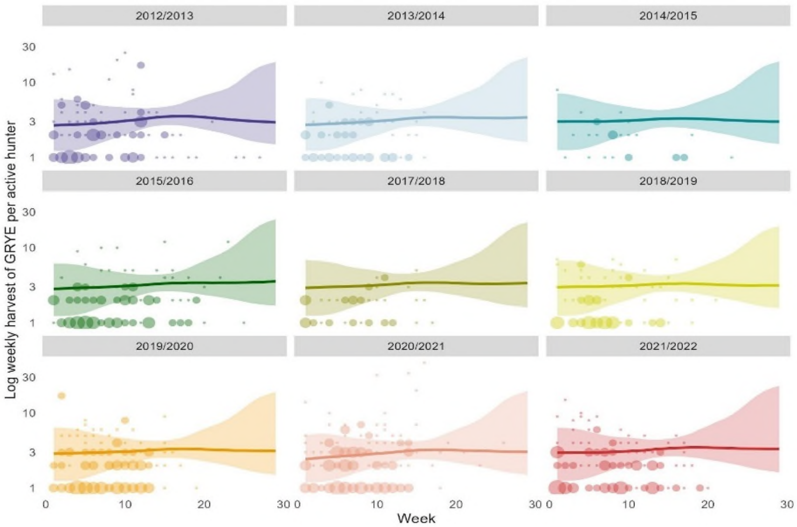


**Figure S6:** Reported weekly harvest of Greater yellowlegs per successful hunter. Solid lines represented median model predicted harvest per hunter. The shaded areas represent the 90% BCIs. Points represent logbook reported harvest, with point size representing the number of hunters reporting the given weekly harvest.

*Pectoral sandpiper*

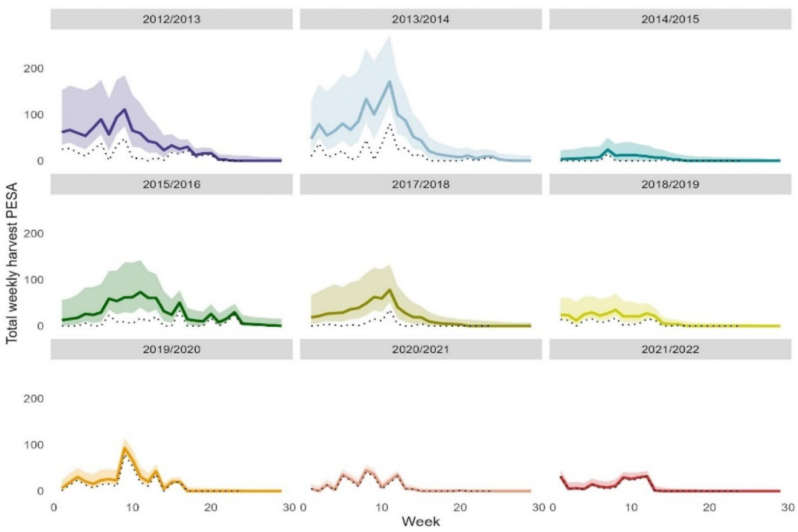


**Figure S7:** Estimated weekly harvest of Pectoral sandpiper. The solid lines represent weekly national harvest using median model predicted weekly harvest, with a shaded area to represent the 90% BCIs on weekly harvest.The dotted line represents the weekly harvest reported in the logbooks.

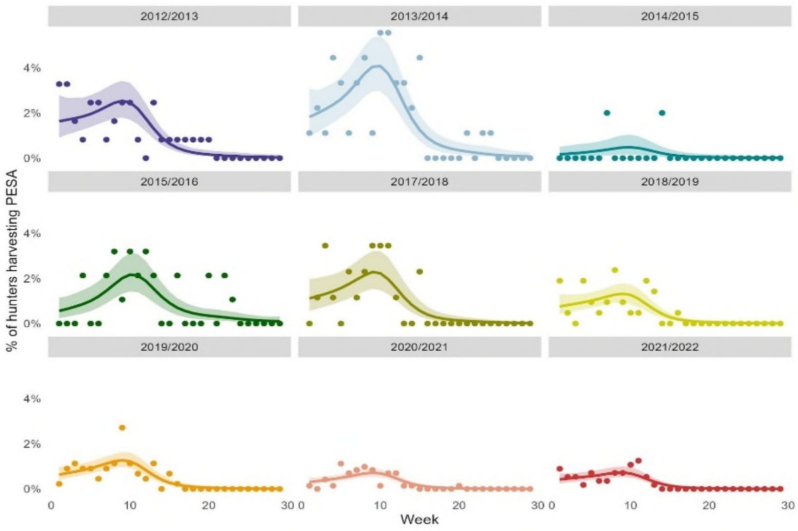


**Figure S8:** Estimated proportion of hunters that successfully harvested Pectoral sandpiper per week. Solid lines represented the median model predicted proportion of hunters harvesting Pectoral sandpipers. The shaded areas represent the 90% BCIs. Points represent the percentage of logbooks in a given week that report harvest of Pectoral sandpipers.

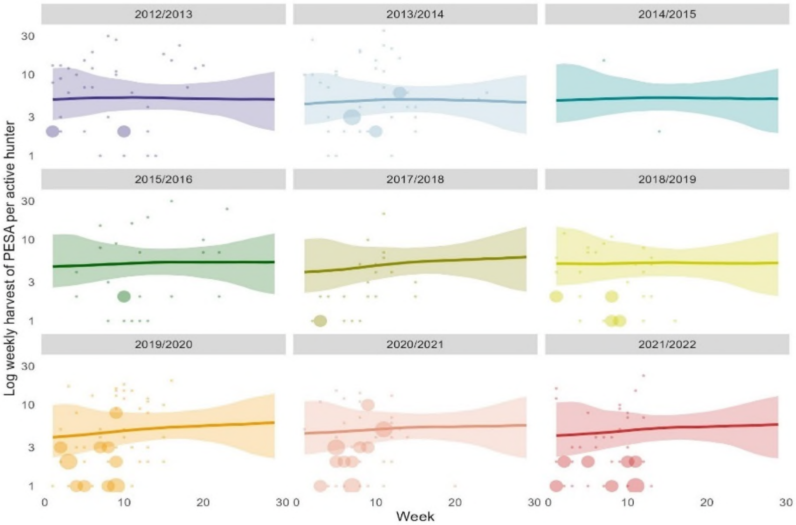


**Figure S9:** Reported weekly harvest of Pectoral sandpiper per successful hunter. Solid lines represented median model predicted harvest per hunter. The shaded areas represent the 90% BCIs. Points represent logbook reported harvest, with point size representing the number of hunters reporting the given weekly harvest.

*Short-billed dowitcher*

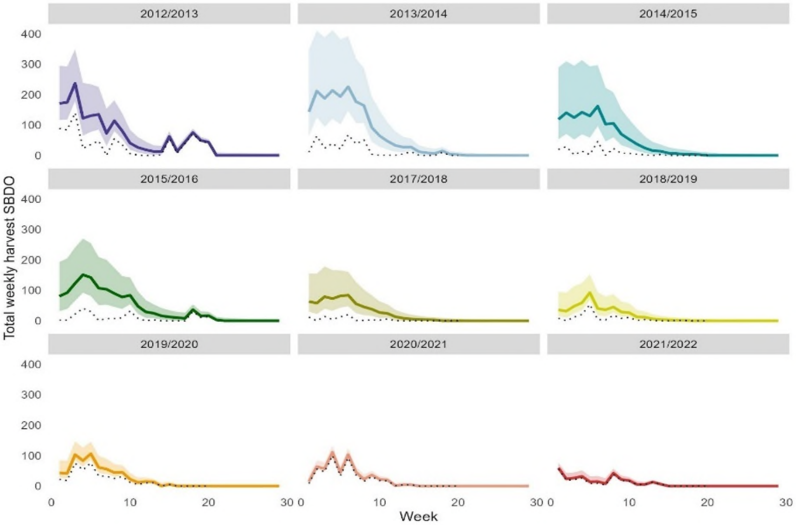


**Figure S10:** Estimated weekly harvest of Short-billed dowitcher. The solid lines represent weekly national harvest using median model predicted weekly harvest, with a shaded area to represent the 90% BCIs on weekly harvest. The dotted line represents the weekly harvest reported in the logbooks.

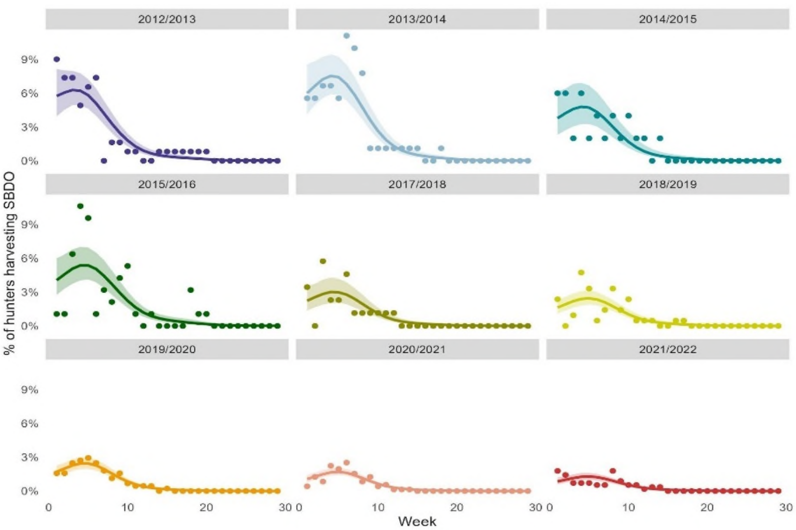


**Figure S11:** Estimated proportion of hunters that successfully harvested Short-billed dowitcher per week. Solid lines represented the median model predicted proportion of hunters harvesting Short-billed dowitchers. The shaded areas represent the 90% BCIs. Points represent the percentage of logbooks in a given week that report harvest of Short-billed dowitchers.

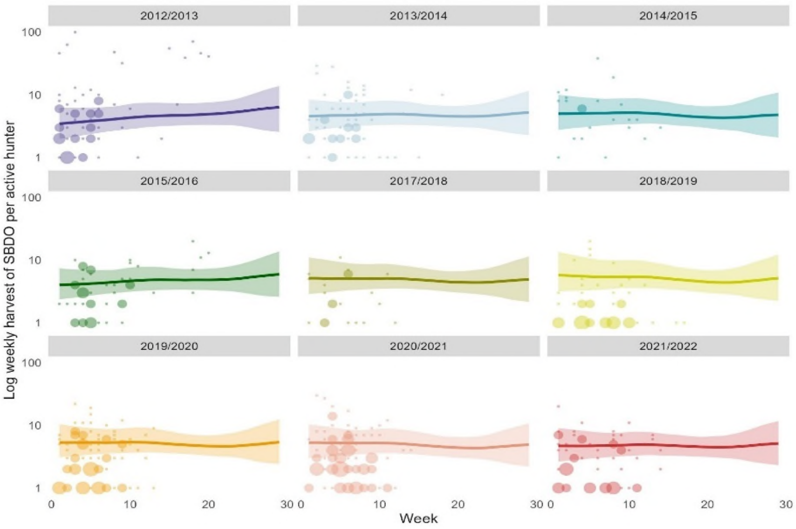


**Figure S12:** Reported weekly harvest of American golden plover per successful hunter. Solid lines represented median model predicted harvest per hunter. The shaded areas represent the 90% BCIs. Points represent logbook reported harvest, with point size representing the number of hunters reporting the given weekly harvest.

*Stilt sandpiper*

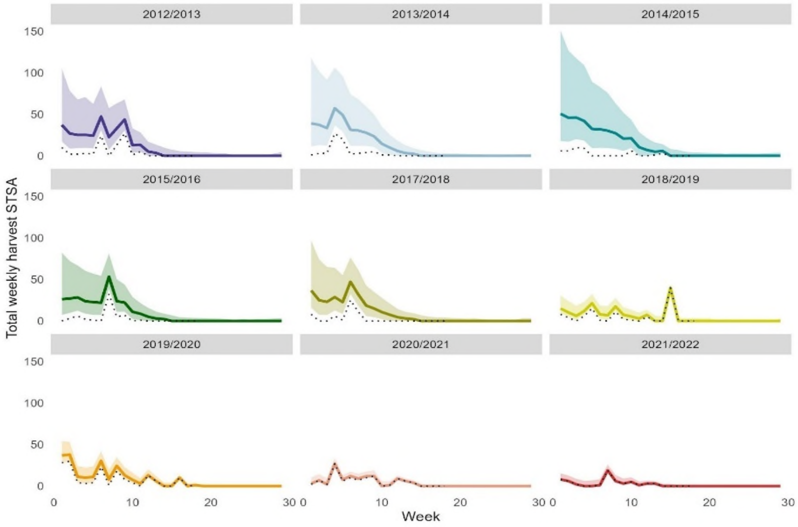


**Figure S13:** Estimated weekly harvest of Stilt sandpiper. The solid lines represent weekly national harvest using median model predicted weekly harvest, with a shaded area to represent the 90% BCIs on weekly harvest.The dotted line represents the weekly harvest reported in the logbooks.

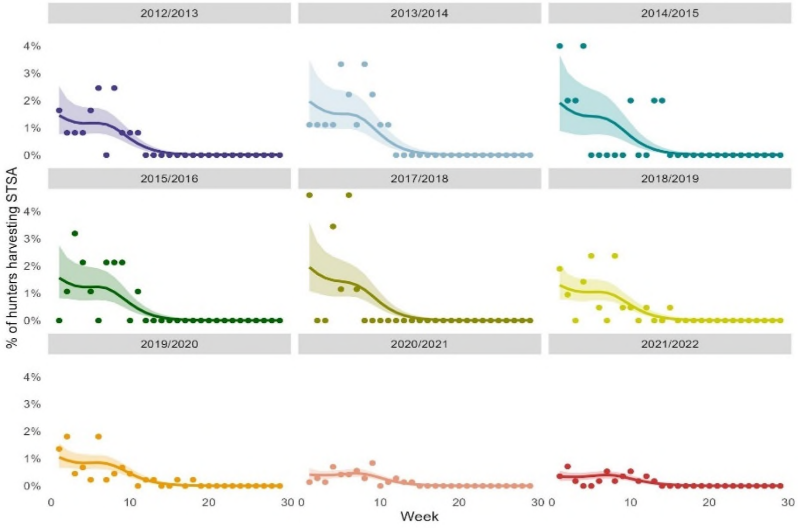


**Figure S14:** Estimated proportion of hunters that successfully harvested Stilt sandpiper per week. Solid lines represented the median model predicted proportion of hunters harvesting Stilt sandpipers. The shaded areas represent the 90% BCIs. Points represent the percentage of logbooks in a given week that report harvest of Stilt sandpipers.

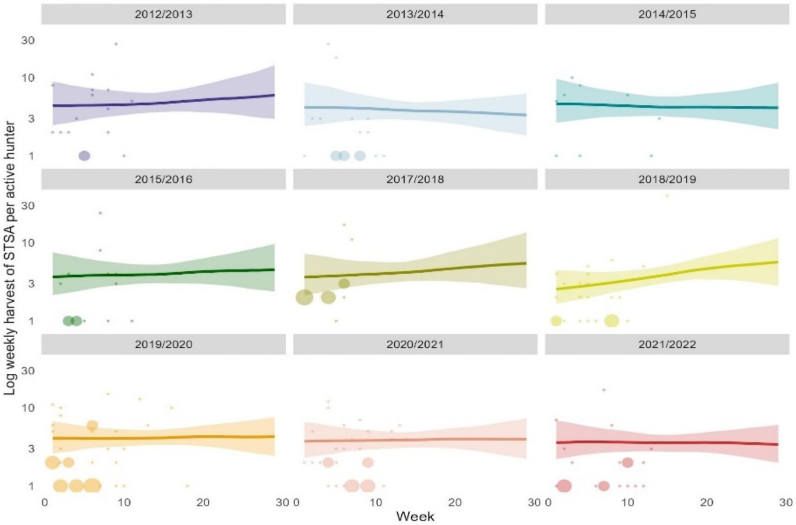


**Figure S15:** Reported weekly harvest of Stilt sandpiper per successful hunter. Solid lines represented median model predicted harvest per hunter. The shaded areas represent the 90% BCIs. Points represent logbook reported harvest, with point size representing the number of hunters reporting the given weekly harvest.

*Teal species*

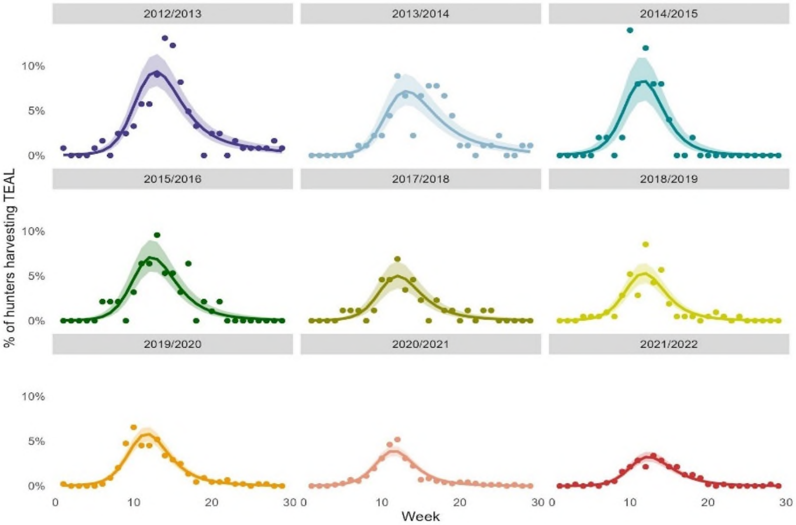


**Figure S16:** Estimated proportion of hunters that successfully harvested teals per week. Solid lines represented the median model predicted proportion of hunters harvesting teals. The shaded areas represent the 90% BCIs. Points represent the percentage of logbooks in a given week that report harvest of teals.

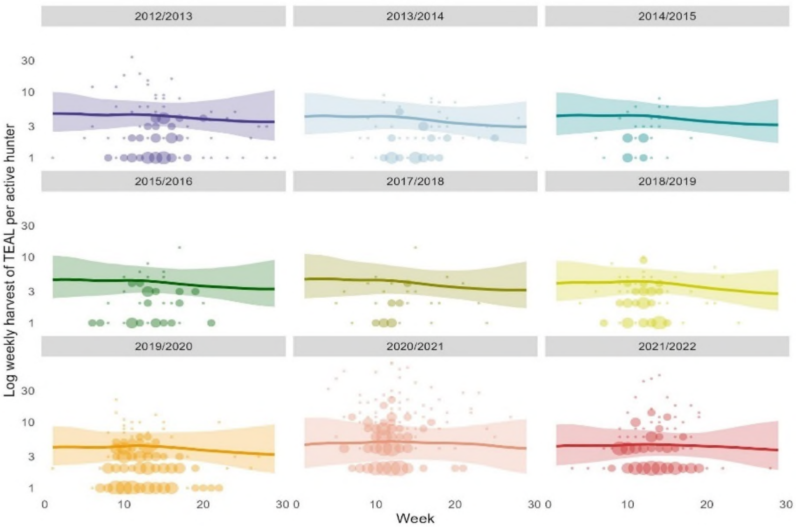


**Figure S17:** Reported weekly harvest of Teal per successful hunter. Solid lines represented median model predicted harvest per hunter. The shaded areas represent the 90% BCIs. Points represent logbook reported harvest, with point size representing the number of hunters reporting the given weekly harvest.

*Upland sandpiper*

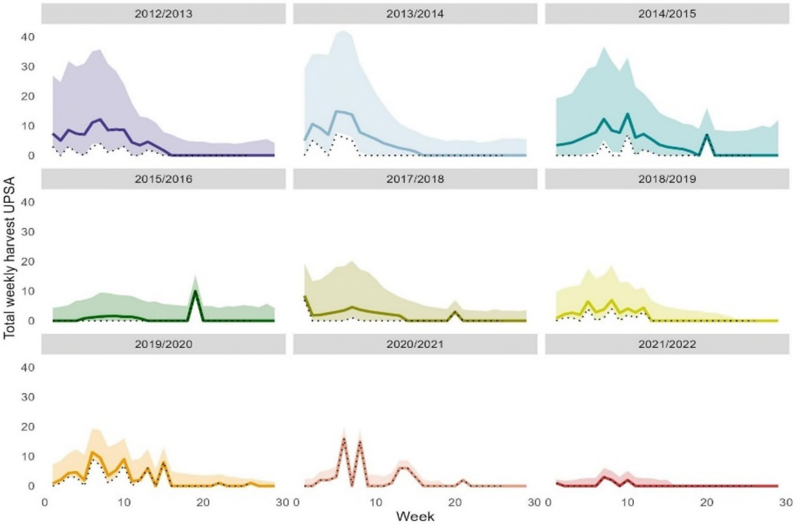


**Figure S18:** Estimated weekly harvest of Upland sandpiper. The solid lines represent weekly national harvest using median model predicted weekly harvest, with a shaded area to represent the 90% BCIs on weekly harvest. The dotted line represents the weekly harvest reported in the logbooks.

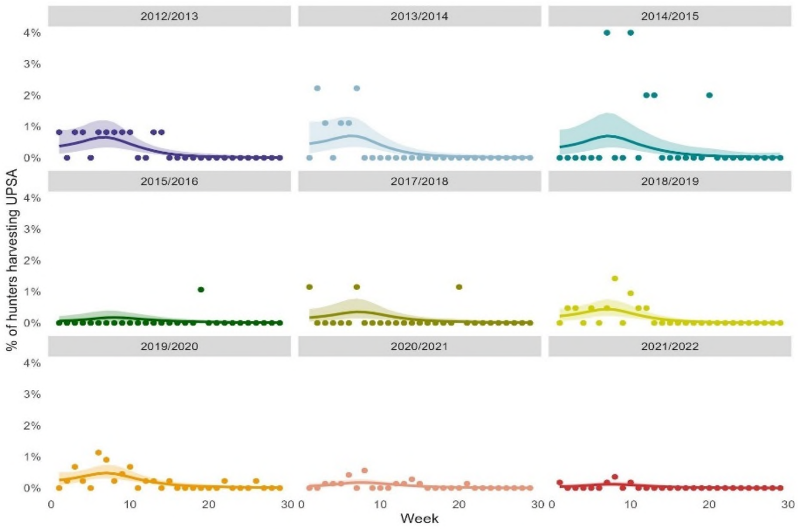


**Figure S19:** Estimated proportion of hunters that successfully harvested Upland sandpiper per week. Solid lines represented the median model predicted proportion of hunters harvesting Upland sandpiper. The shaded areas represent the 90% BCIs. Points represent the percentage of logbooks in a given week that report harvest of Upland sandpiper.

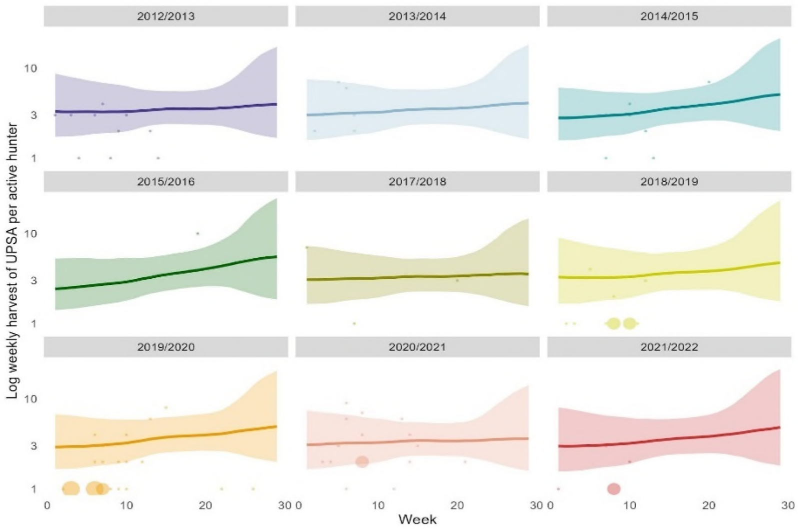


**Figure S20:** Reported weekly harvest of Upland sandpiper per successful hunter. Solid lines represented median model predicted harvest per hunter. The shaded areas represent the 90% BCIs. Points represent logbook reported harvest, with point size representing the number of hunters reporting the given weekly harvest.

*Whimbrel*

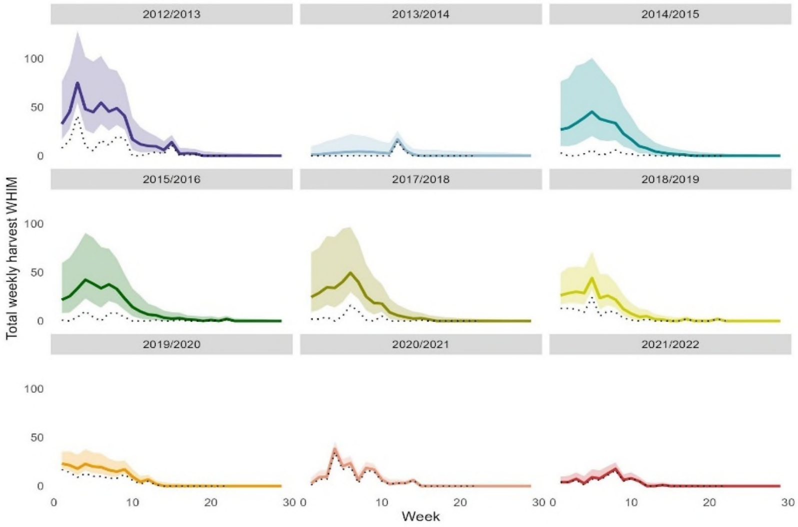


**Figure S21:** Estimated weekly harvest of Whimbrel. The solid lines represent weekly national harvest using median model predicted weekly harvest, with a shaded area to represent the 90% BCIs on weekly harvest.The dotted line represents the weekly harvest reported in the logbooks.

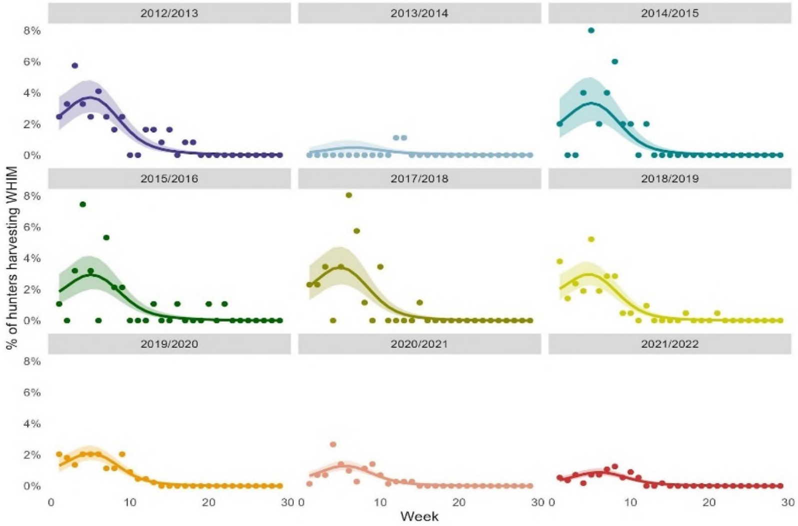


**Figure S22:** Estimated proportion of hunters that successfully harvested Whimbrels per week. Solid lines represented the median model predicted proportion of hunters harvesting Whimbrels. The shaded areas represent the 90% BCIs. Points represent the percentage of logbooks in a given week that report harvest of Whimbrels.

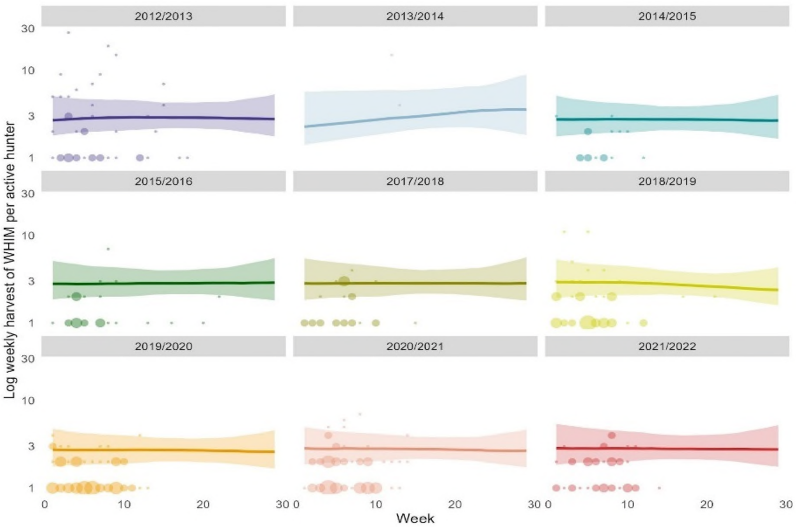


**Figure S23:** Reported weekly harvest of Whimbrel per successful hunter. Solid lines represented median model predicted harvest per hunter. The shaded areas represent the 90% BCIs. Points represent logbook reported harvest, with point size representing the number of hunters reporting the given weekly harvest.

*Willet*

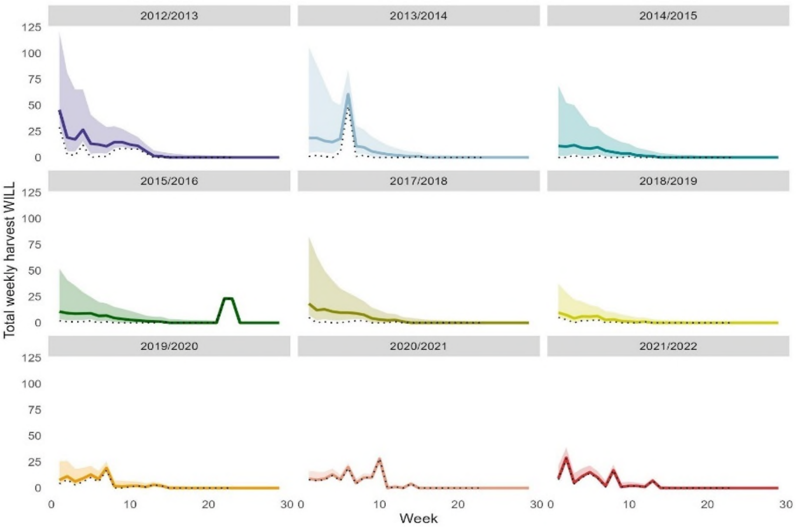


**Figure S24:** Estimated weekly harvest of Willet. The solid lines represent weekly national harvest using median model predicted weekly harvest, with a shaded area to represent the 90% BCIs on weekly harvest. The dotted line represents the weekly harvest reported in the logbooks.

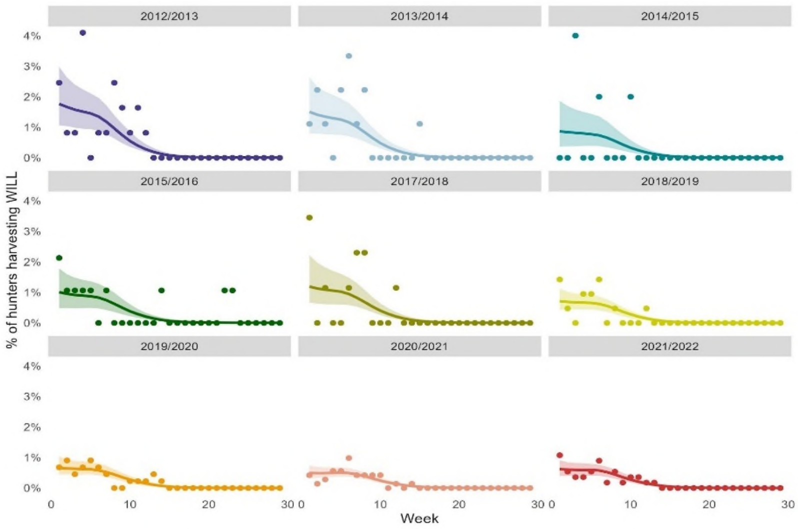


**Figure S25:** Estimated proportion of hunters that successfully harvested Willets per week. Solid lines represented the median model predicted proportion of hunters harvesting Willets. The shaded areas represent the 90% BCIs. Points represent the percentage of logbooks in a given week that report harvest of Willets.

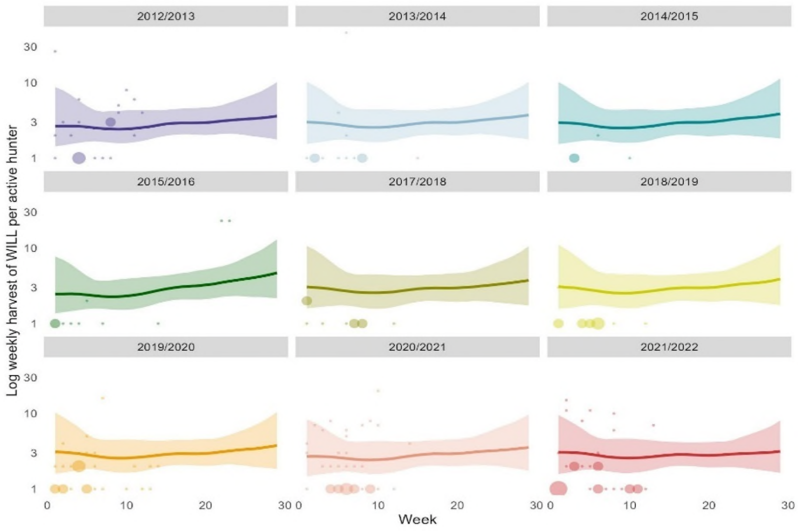


**Figure S26:** Reported weekly harvest of Willet per successful hunter. Solid lines represented median model predicted harvest per hunter. The shaded areas represent the 90% BCIs. Points represent logbook reported harvest, with point size representing the number of hunters reporting the given weekly harvest.

**Resident species**

*Common ground dove*

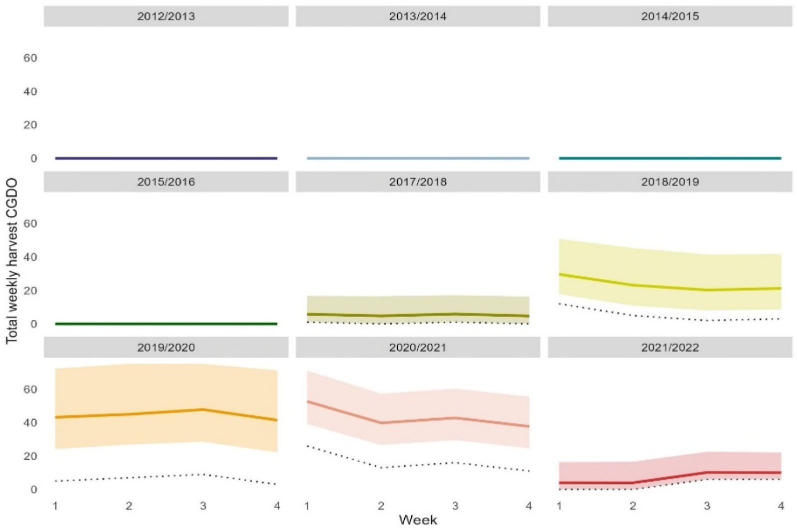


**Figure S27:** Estimated weekly harvest of Common ground dove. The solid lines represent weekly national harvest using median model predicted weekly harvest, with a shaded area to represent the 90% BCIs on weekly harvest.The dotted line represents the weekly harvest reported in the logbooks.

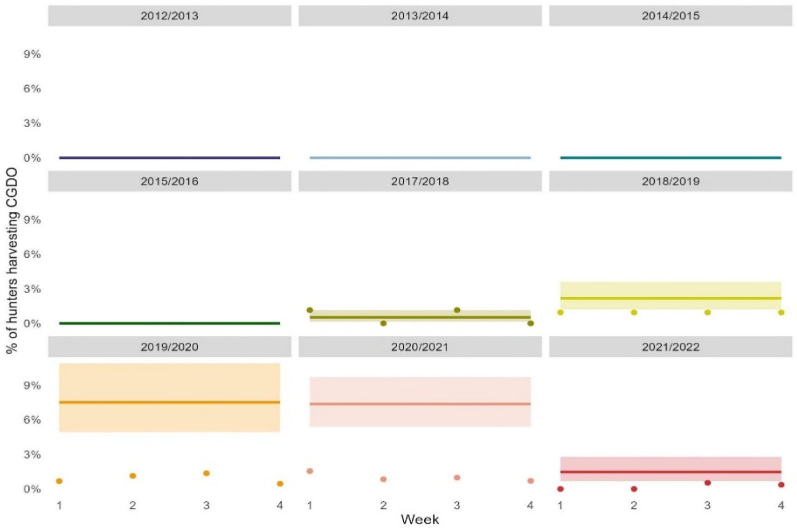


**Figure S28**: Estimated proportion of hunters that successfully harvested Common ground doves per week. Solid lines represented the median model predicted proportion of hunters harvesting Common ground doves. The shaded areas represent the 90% BCIs. Points represent the percentage of logbooks in a given week that report harvest of Common ground doves.

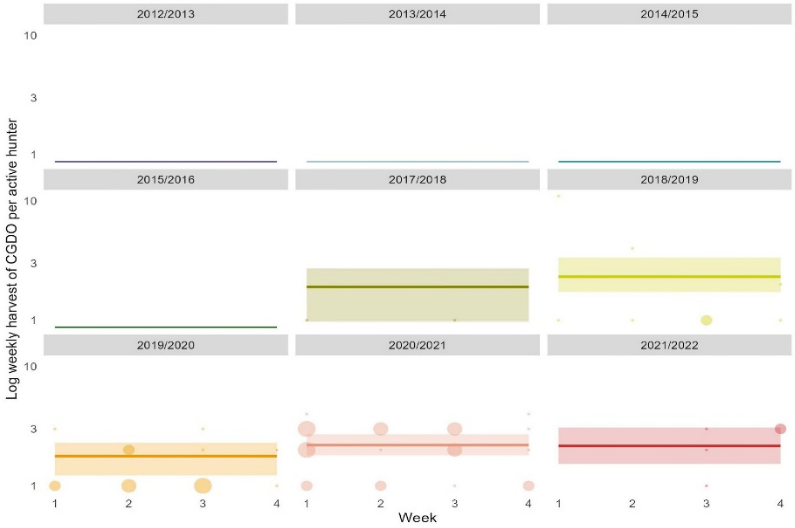


**Figure S29:** Reported weekly harvest of Common ground dove per successful hunter. Solid lines represented median model predicted harvest per hunter. The shaded areas represent the 90% BCIs. Points represent logbook reported harvest, with point size representing the number of hunters reporting the given weekly harvest.

*Eurasian Collared Dove*

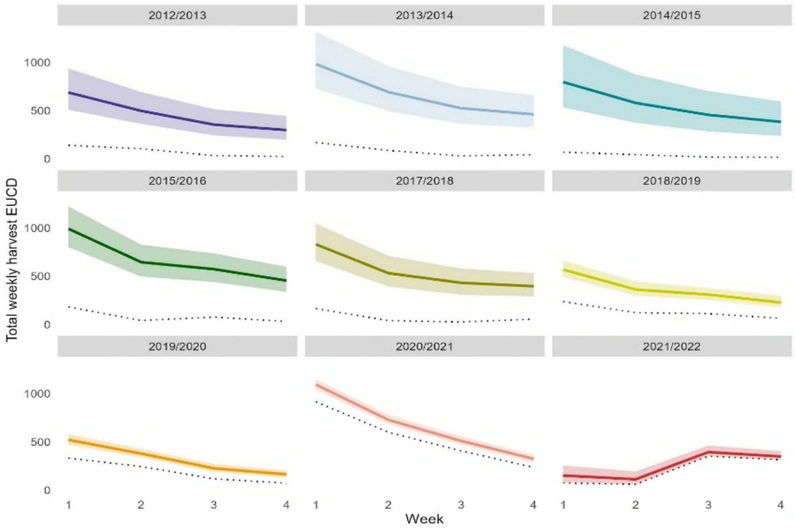


**Figure S30:** Estimated weekly harvest of Eurasian collared dove. The solid lines represent weekly national harvest using median model predicted weekly harvest, with a shaded area to represent the 90% BCIs on weekly harvest.The dotted line represents the weekly harvest reported in the logbooks.

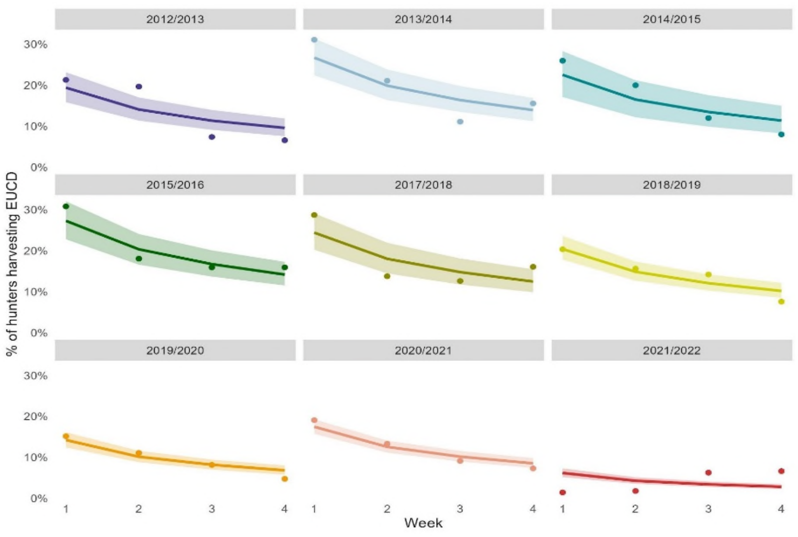


**Figure S31**: Estimated proportion of hunters that successfully harvested Eurasian collared doves per week. Solid lines represented the median model predicted proportion of hunters harvesting Eurasian collared doves. The shaded areas represent the 90% BCIs. Points represent the percentage of logbooks in a given week that report harvest of Eurasian collared doves.

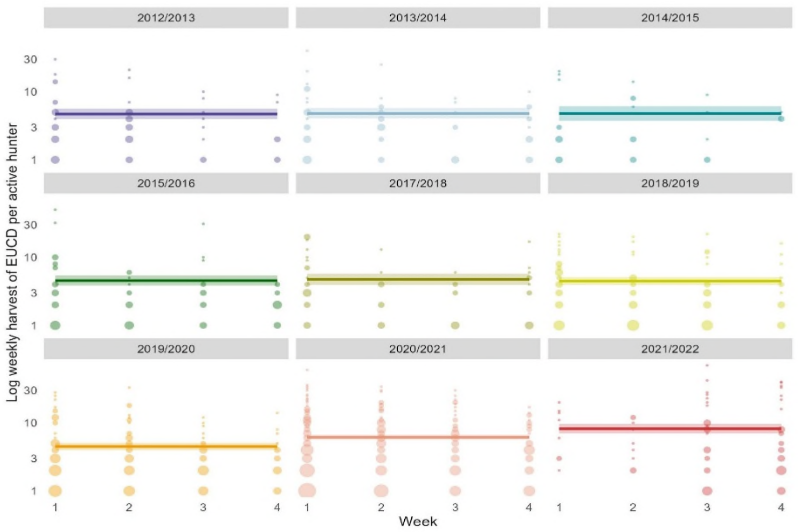


**Figure S32:** Reported weekly harvest of Eurasian collared dove per successful hunter. Solid lines represented median model predicted harvest per hunter. The shaded areas represent the 90% BCIs. Points represent logbook reported harvest, with point size representing the number of hunters reporting the given weekly harvest.

*Pearly-eyed thrasher*

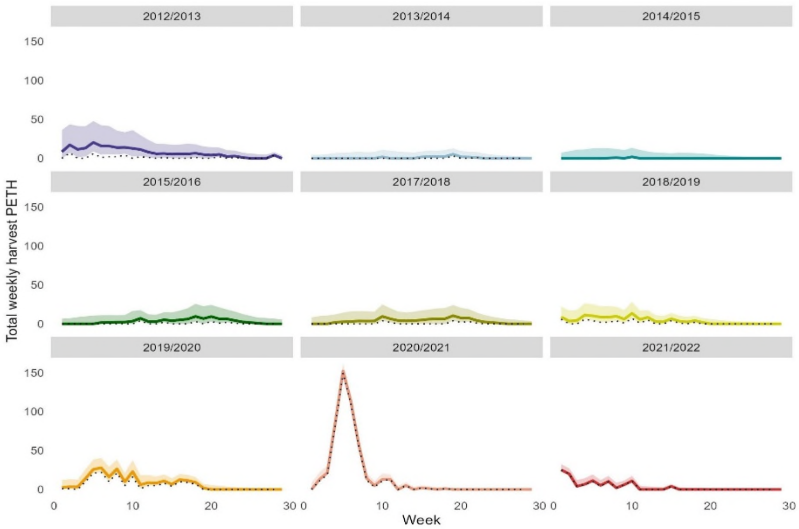


**Figure S33:** Estimated weekly harvest of Pearly-eyed thrasher. The solid lines represent weekly national harvest using median model predicted weekly harvest, with a shaded area to represent the 90% BCIs on weekly harvest.The dotted line represents the weekly harvest reported in the logbooks.

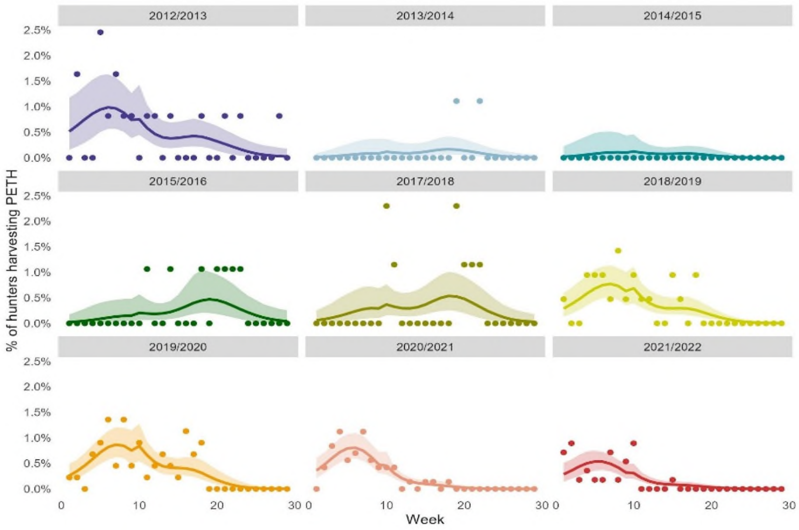


**Figure S34**: Estimated proportion of hunters that successfully harvested Pearly-eyed thrasher per week. Solid lines represented the median model predicted proportion of hunters harvesting Pearly-eyed thrasher. The shaded areas represent the 90% BCIs. Points represent the percentage of logbooks in a given week that report harvest of Pearly-eyed thrasher doves.

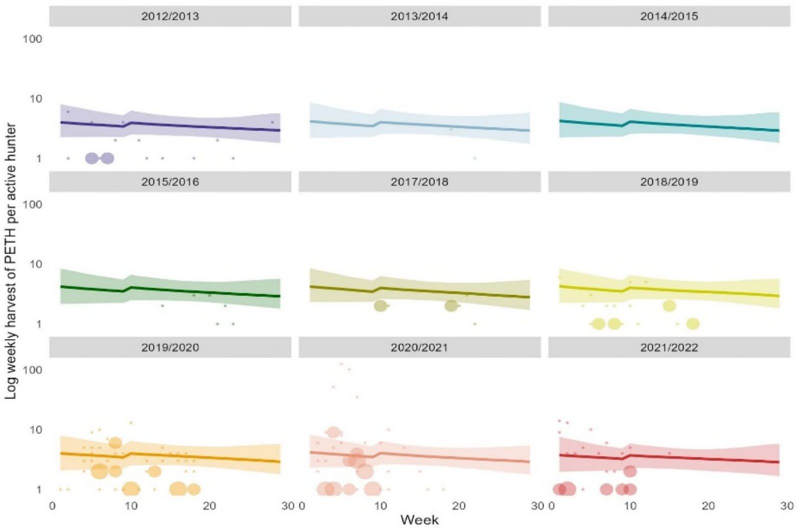


**Figure S35:** Reported weekly harvest of pearly-eyed thrasher per successful hunter. Solid lines represented median model predicted harvest per hunter. The shaded areas represent the 90% BCIs. Points represent logbook reported harvest, with point size representing the number of hunters reporting the given weekly harvest.

Scaly-breasted thrasher

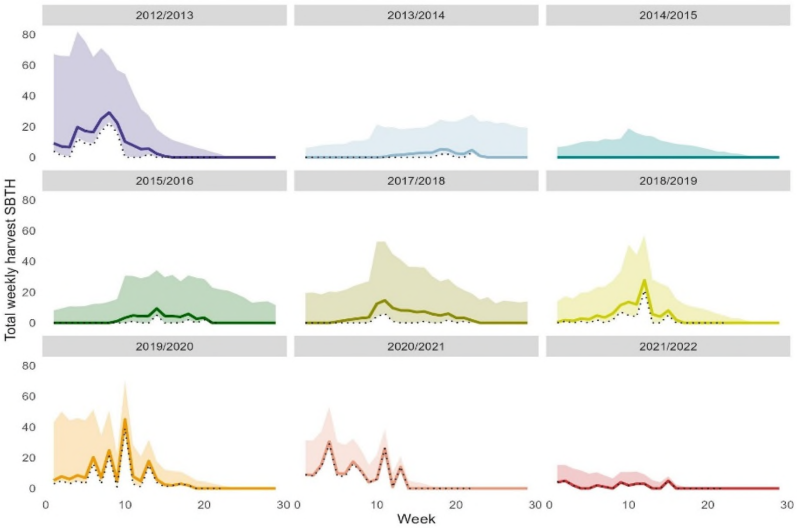


**Figure S36:** Estimated weekly harvest of Scaly-breasted thrasher. The solid lines represent weekly national harvest using median model predicted weekly harvest, with a shaded area to represent the 90% BCIs on weekly harvest.The dotted line represents the weekly harvest reported in the logbooks.

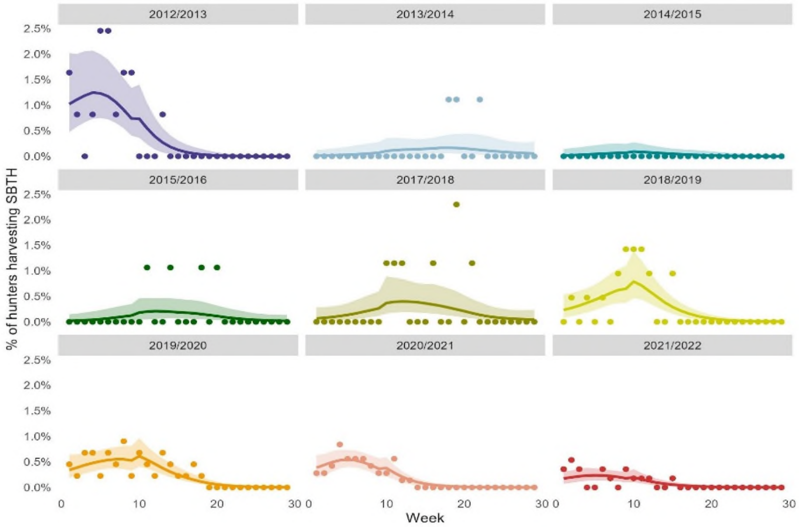


**Figure S37**: Estimated proportion of hunters that successfully harvested Scaly-breasted thrasher per week. Solid lines represented the median model predicted proportion of hunters harvesting Scaly breasted thrasher. The shaded areas represent the 90% BCIs. Points represent the percentage of logbooks in a given week that report harvest of Scaly-breasted thrasher.

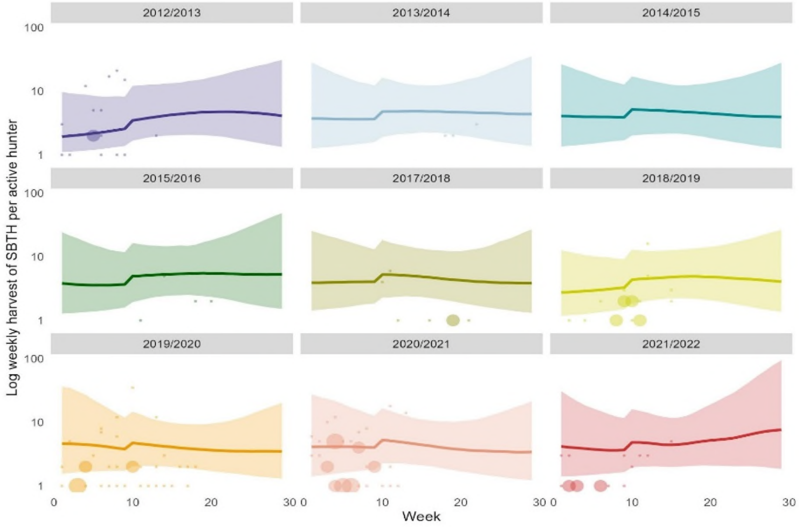


**Figure S38: *Reported weekly harvest of Scaly-breasted thrasher per successful hunter. Solid lines represented median model predicted harvest per hunter. The shaded areas represent the 90% BCIs. Points represent logbook reported harvest, with point size representing the number of hunters reporting the given weekly harvest.

*Scaly-naped pigeon*

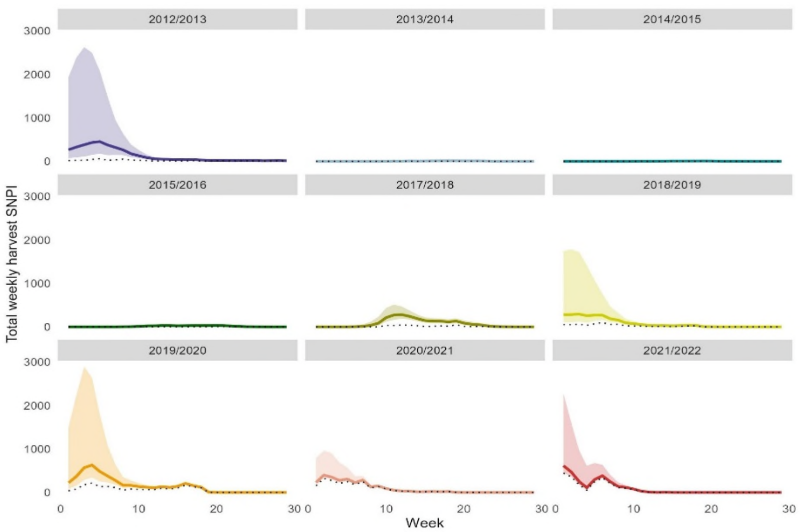


**Figure S39:** Estimated weekly harvest of Scaly-naped pigeon. The solid lines represent weekly national harvest using median model predicted weekly harvest, with a shaded area to represent the 90% BCIs on weekly harvest.The dotted line represents the weekly harvest reported in the logbooks.

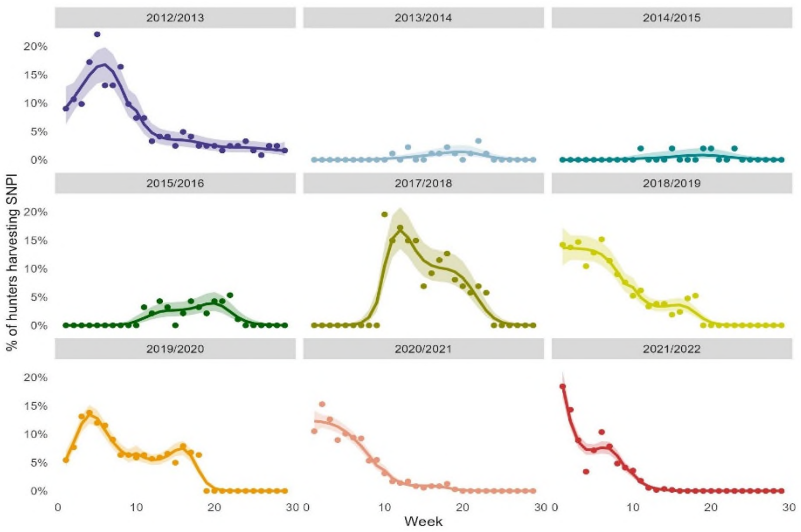


**Figure S40**: Estimated proportion of hunters that successfully harvested Scaly-naped pigeon per week. Solid lines represented the median model predicted proportion of hunters harvesting Scaly-naped pigeon. The shaded areas represent the 90% BCIs. Points represent the percentage of logbooks in a given week that report harvest of Scaly-naped pigeon.

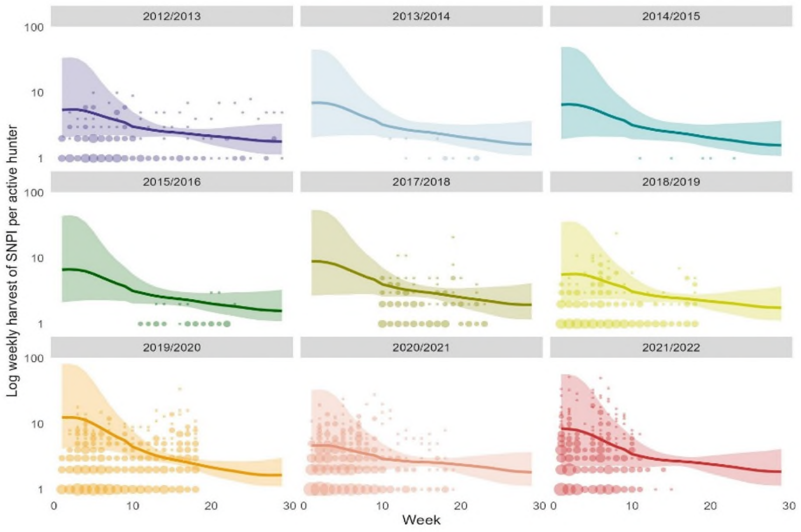


**Figure S41:** Reported weekly harvest of Scaly-naped pigeon per successful hunter. Solid lines represented median model predicted harvest per hunter. The shaded areas represent the 90% BCIs. Points represent logbook reported harvest, with point size representing the number of hunters reporting the given weekly harvest.

White-crowned pigeon

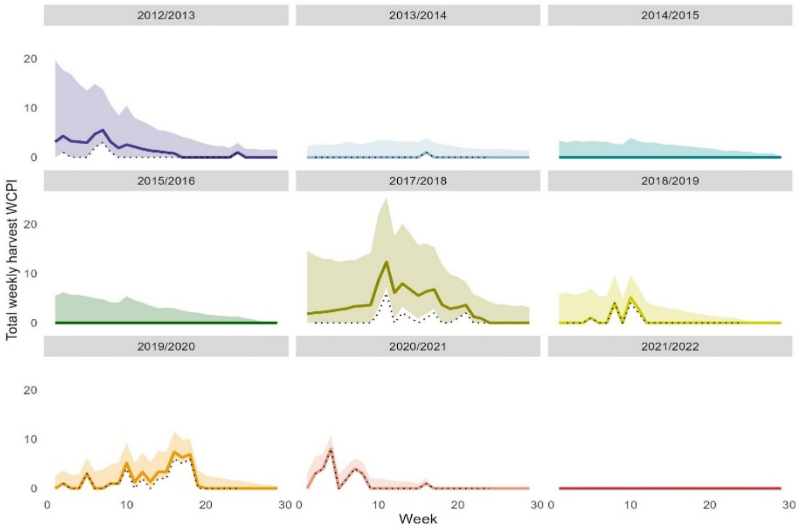


**Figure S42:** Estimated weekly harvest of White-crowned pigeon. The solid lines represent weekly national harvest using median model predicted weekly harvest, with a shaded area to represent the 90% BCIs on weekly harvest.The dotted line represents the weekly harvest reported in the logbooks.

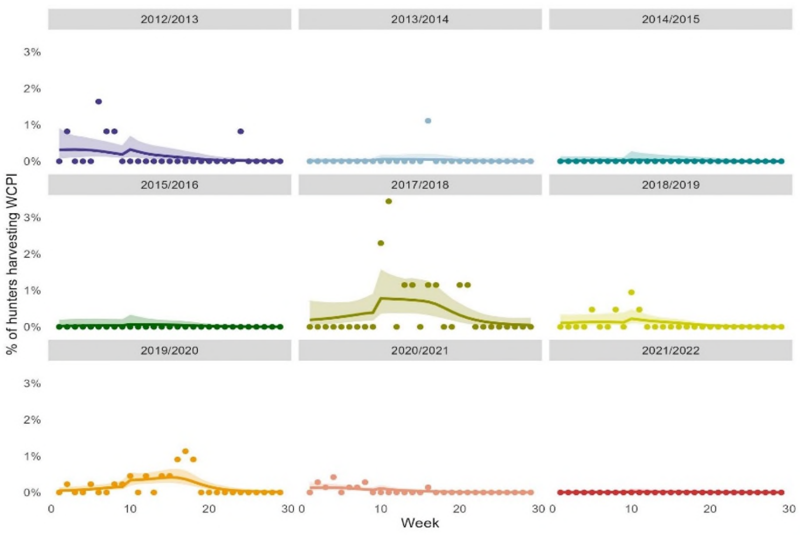


**Figure S43**: Estimated proportion of hunters that successfully harvested White-crowned pigeon per week. Solid lines represented the median model predicted proportion of hunters harvesting White-crowned pigeon. The shaded areas represent the 90% BCIs. Points represent the percentage of logbooks in a given week that report harvest of White-crowned pigeon.

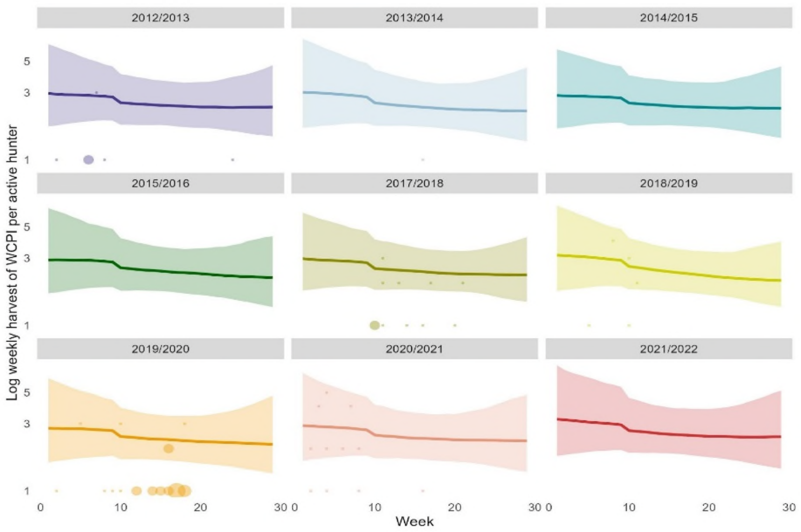


**Figure S44:** Reported weekly harvest of White-crowned pigeon per successful hunter. Solid lines represented median model predicted harvest per hunter. The shaded areas represent the 90% BCIs. Points represent logbook reported harvest, with point size representing the number of hunters reporting the given weekly harvest.

*Zenaida dove*

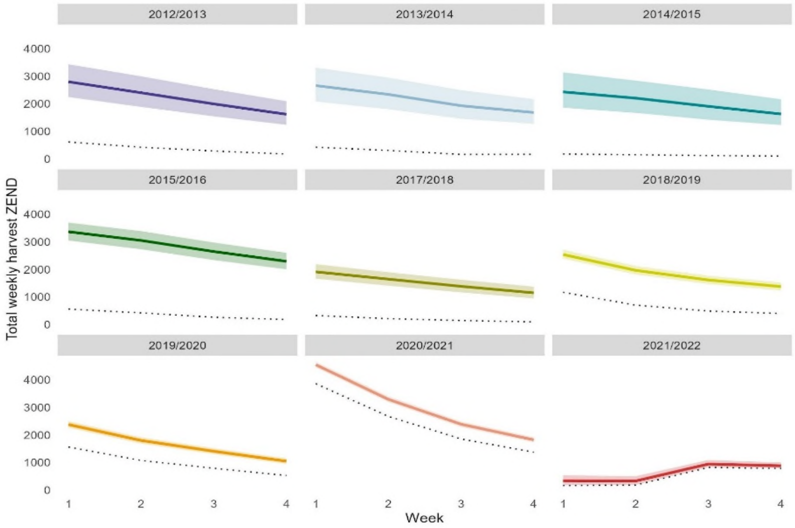


**Figure S45:** Estimated weekly harvest of Zenaida dove. The solid lines represent weekly national harvest using median model predicted weekly harvest, with a shaded area to represent the 90% BCIs on weekly harvest.The dotted line represents the weekly harvest reported in the logbooks.

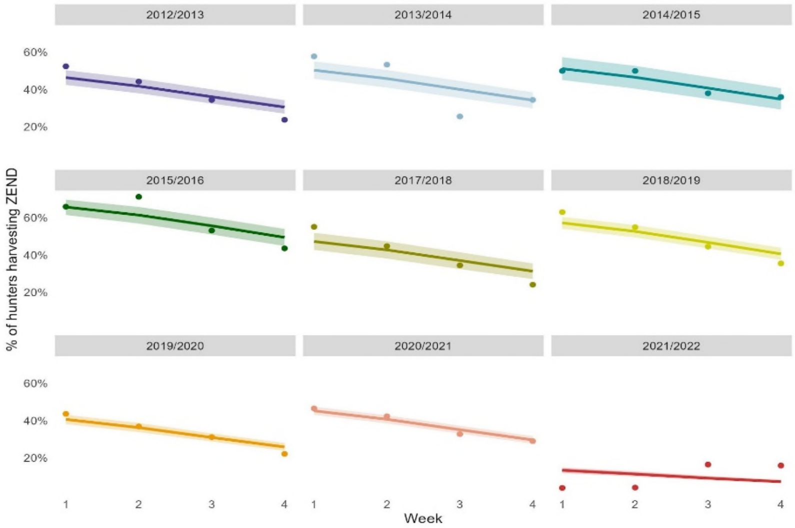


**Figure S46**: Estimated proportion of hunters that successfully harvested Zenaida doves per week. Solid lines represented the median model predicted proportion of hunters harvesting Zenaida doves. The shaded areas represent the 90% BCIs. Points represent the percentage of logbooks in a given week that report harvest of Zenaida doves.

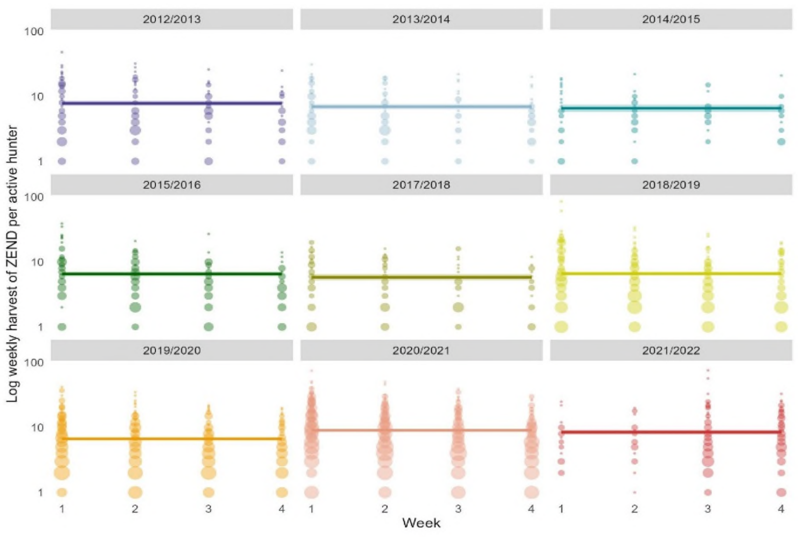


**Figure S47:** Reported weekly harvest of Zenaida dove per successful hunter. Solid lines represented median model predicted harvest per hunter. The shaded areas represent the 90% BCIs. Points represent logbook reported harvest, with point size representing the number of hunters reporting the given weekly harvest.
